# Competing endogenous RNA network: Potential entrants to gene editing in Hepatocellular Carcinoma Gene editing of ceRNA in Hepatocellular Carcinoma

**DOI:** 10.1101/288381

**Authors:** Ayman El-Sayed Shafei, Marwa Matboli, Mahmoud A. Ali, Ziad Nagy, Maged Reda, Mohamed Salah, Ahmed Hamdy, Mahmoud Abdelgawad, Ahmed Ashry, Mohammad Tarek, Osama Saber, Ahmed Azazy, Badr Mohamed, Mohmed K. Hassan, Nashwa El-Khazragy, Sherif El-Khamisy

## Abstract

**Background:** Hepatocellular Carcinoma (HCC) is the leading cause of cancer deaths worldwide as well as in Egypt. We aimed to use Clustered Regulatory Interspaced Short Palindromic Repeats (CRISPR) gene editing technique to induce forced down-regulation of the circRNA which consequently modified miRNA expression in HepG2 cell line to prove the regulatory relationship between the RNA parts of an in silico-detected competing endogenous RNA network in HCC

**Method:** We first retrieved hsa_circ_0000064-miR-1285-TRIM2 mRNA from public microarray databases followed by in silico modelling to mimic the regulation kinetics of cirRNA associated ceRNA network. Secondly, we performed polymerase chain reaction (PCR)-based amplification of synthetic fragments, Gibson assembly of both CRISPR and non CRISPR based circuits, E-coli transformation, plasmid purification, HePG2 cell line transfection. Finally Expression levels of the chosen RNAs in hepatocellular carcinoma (HCC) cell line, HepG2, were examined by quantitative reverse transcription polymerase chain reaction (qRT-PCR) and the cytotoxic effect was validated by viability assay.TRIM2 protein expression was proved by immunohistochemistry and flowcytometry.

**Results:** Induction of hsa_circ_0000064 into HepG2 cell line via CRISPR-and non-CRISPR mediated synthetic circuit resulted in statistically significant decrease in cell number and, then, cellular viability with marked increase in hsa_circ_0000064 and TRIM2 mRNA levels and concomitant decrease in miR-1285 expression in HepG2 cell line compared with control (p<0.0). Moreover exogenous expression of hsa_circ_0000064 in HepG2 cell line showed increased expression of the tumor suppressor protein, TRIM2.

**Conclusions:** Our integrative approach, including in silico data analysis and experimental validation proved that CRISPR-mediated synthetic circuit-based overexpression of hsa_circ_0000064 was more efficient than conventional transient transfection, representing a promising therapeutic strategy for HCC.

**Data Availability:** Our Data was made available online on the IGEM wiki of team AFCM-EGYPT: http://2017.igem.org/Team:AFCM-Egypt. Synthetic parts have been submitted to IGEM Parts Registry.

**Financial Disclosure:** The project was funded by Armed Forces College of Medicine AFCM, Zewail City of Science and Technology, National Research Center NRC, VitaBiotics, PHARCO Pharmaceuticals, Sim Era and DANUB Paintings. IDT provided 20 kb of DNA synthesis. The funders had no role in study design, data collection and analysis, decision to publish, or preparation of the manuscript.

## Introduction

Liver cancer is one of the leading causes of cancer deaths worldwide, accounting for more than 600,000 deaths each year. In the united states alone, it’s estimated that there are about 39,230 of newly diagnosed cases and about 27,170 reported deaths of primary liver cancer and intrahepatic bile duct cancers in 2016 [1]. In Egypt, liver cancer is a serious if not the most serious cancer problem. It is ranked the first among cancers in males (33.6%) and equally with breast cancer among females based upon results of the National Cancer Registry Program (NCRP 2008-2011) [2]. The rising rates of HCC in Egypt are due to the high prevalence of hepatitis B virus (HBV) and hepatitis C virus infection (HCV) among Egyptian population [3]. According to literature, El-Zayadi et al. reported almost 2 folds increase in HCC among chronic liver disease patients over a decade [4]. Also, according to Ibrahim et al, HCC is the first most common cancer in males and second most common cancer in females [2]. Therefore, we need effective strategies for early detection and better management of HCC which will be of great value in developing countries with limited resources and high incidence rates of HCC.

It is well known that the human genome is actively transcribed. However, there are only about 20,000 protein-coding genes, accounting for about 2% of the genome and the rest of the transcripts are non-coding RNAs including microRNAs and long non-coding RNAs (lncRNAs). Non-coding RNAs play an important role in the regulation of gene expression, including chromatin modification, transcription and post-transcriptional processing. It has been confirmed that dysregulation of non-coding RNAs is accompanied by a number of human pathological diseases, mainly tumors [5]. The interplay between diverse RNA species, whereby one transcript can reciprocally modulate the expression of another transcript by sequestering shared miRNAs, has been referred to as ceRNA crosstalk. CeRNA activity has been attributed to both protein-coding and non-coding RNA transcripts. Although ceRNA research is rapidly growing, accumulating evidence suggests that this additional dimension of post-transcriptional gene regulation, in which RNA transcripts titrate miRNA availability, represents a biologically relevant, well-conserved and widespread mechanism of regulation [6]. CircRNAs are a large class of RNAs that have shown huge capability as gene regulators in humans. Beyond being a potentially major approach of gene regulation, circRNAs may represent new roles in cancer diagnosis and targeted therapy. CircRNAs have been previously shown to play a role in the development and progression of HCC. Thereby, modulation of cirRNA and miRNA by gene editing tools may significantly decrease cell viability, inhibit motility and invasiveness and sensitize cells to multi-stimuli-induced apoptosis [7–9].

Circular RNAs have been reported to act in a sponge manner regulating miRNAs in competing endogenous RNA networks (ceRNA) which presents a promising role regarding their therapeutic and diagnostic potential. Circular RNA has_circ_0004277 have been reported to possess potential diagnostic implications for acute myeloid leukemia [10]. Circular RNAs has_circ_0013958 and has_circ_0000190 have been also reported to have of promising diagnostic value for lung cancer and gastric cancer respectively [11]. Other circular RNAs were reported to be of therapeutic importance including has_circ_0016347 which regulates invasion and proliferation of osteosarcoma [12]. These circular RNAs usually target mRNAs through regulating miRNAs, so for example circular RNA has_circ_0045714 have been proved to apoptotic pathways of chondrocytes through targeting miR-193b/IGF1R axis [13]. In a study conducted on colorectal cancer tissues, hsa_circ_001569 was selected as a potential regulator of cancer progression, with high expression level detected by RT-qPCR analysis. Whereas the interaction with miR-145, miR-145 was found to target the 3′ UTR of E2F5, BAG4 and FMNL2 transcripts in colorectal cells, reducing their mRNA expression levels, however the presence of circ_001569 increased the protein levels of these transcripts where it was also reported that disrubitng expression of circRNA resulted in deregulation of target gene expression through miRNA-mediated circRNA-associated ceRNA crosstalk interactions [14].

Many attempts have been made to restore the function of dysfunctional genes by gene knock-down using small interfering RNAs (siRNAs) and microRNAs (miRNAs) [15]. There are several genetic tools available for this purpose, including zinc finger nuclease (ZFN) and transcription activation like element nuclease (TALEN) [16] Recently, a novel genetic engineering tool called clustered regularly interspaced short palindromic repeats (CRISPR)/CRISPR-associated (Cas) system is more advanced because of easy generation and high efficiency of gene targeting. Importantly, it only requires changing the sequence of the guide RNA (gRNA). CRISPR/Cas9 has rapidly gained popularity due to its superior simplicity [17]. In this system, a single guide RNA (sgRNA) complexes with Cas9 nuclease, which can recognize a variable 20-nucleotide target sequence adjacent to a 5′-NGG-3′ protospacer adjacent motif (PAM) and introduce a DSB in the target DNA [18]. The induced DSB (DNA double stranded break) then triggers DNA repair process mainly via two distinct mechanisms: namely, the non-homologous end joining (NHEJ) and the homology-directed repair (HDR) pathways. The CRISPR system is meant to provide adaptive immunity against phages and other mobile genetic elements in bacteria and archaea. While most of the early work has largely been dominated by examples of CRISPR-Cas systems directing the cleavage of phage or plasmid [19–21].

In this IGEM project we aimed to analyze circRNA and disease bioinformatics databases to select significantly relevant circRNA for HCC. Furthermore, we aimed to analyze circRNA-miRNA interaction databases to retrieve competing endogenous RNA specific for HCC, with further characterization of the expression of the serum cirRNA-associated ceRNA genes in HepG2 cell line to evaluate their role in pathogenesis of HCC. Finally we wanted to compare between the efficacy of a CRISPR and non-CRISPR based synthetic circuit on modulating cirRNA-associated ceRNA related HCC expression using HepG2 cell line.

## Methods

### Dry Lab Methods

#### Bioinformatics Analysis for Biomarker filtration and Target Determination

We started by filtering HCC-associated circular-RNAs from bioinformatics databases such as Circ2Trait [22] then, we used circInteractome database [23] to retrieve filter the highly associated miRNAs with our Circular RNA on Interest besides determining their binding sites on circular RNA transcript. TargetScan [24] Database bioinformatics tool was used to filter out the associated mRNAs to be regulated in previously Designed ceRNA network.

#### Deterministic Modeling

We have used sysBio [25] R package to simulate models using a differential equation solver. Modeling Protocol for each model started by using new Model function to create a new model. Then we created a model object list specifying information about model including; name, reactions, species, rates, parameters, rules, models, ODEs. Using the addMAreaction function, we added specified reactions into the model - to be interpreted using the law of mass action. We used addMAreactRate and addParameters functions to specify information about the reaction rates and parameters involved in the model. Finally, we defined species using the add Species function. Consequently, we used makeModel function to create a mathematical representation of the model. This function transforms reactions into corresponding ODEs, and creates stochastic matrix and propensity function to perform stochastic modeling. We used simulateModel function to run simulation (solve ODEs). This function calls the validateModel function that checks if all components of the models have been defined. Finally, we used plotResults function to visualize simulation results. The modeling environment has been designed and implemented in an open-source publicly available online tool [26].

#### Stochastic Modeling

Stochasticity is a key player in regulation of gene expression especially when the number of molecular species involved is small. Thus, genetic circuits such as ceRNA networks are usually embedded in more complex networks such as miRNA-Target networks, so that induced interactions might be of regulatory value. A stochastic analysis of the ceRNA system is necessary as potential crosstalk between miRNA targets is quite indicative of the degree of interaction between regulatory networks [27].

##### Circular RNA (hsa_circ_0000064) Structural Modelling

We have used Vienna RNA package [28] for generating structural models of circular RNA hsa_circ_0000064. RNA sequence was retrieved from circInteractome database [23] to be used as an input for Vienna package. Vienna RNA package depends on extension of linear folding algorithms. Circular RNA molecules are modelled through post-processing of computed linear arrays. Using Vienna RNA Package, we could compare structural modifications between linear and circular structures in a memory-effective manner [29]. The energy contribution of Exterior loop should be scored in circular structures, on the other hand, exterior loops have no energy contribution in linear structures. RNAfold structure prediction tool was used to calculate the minimum free energy (MFE) and back traces an optimal secondary structure, mountain plot and dot plots were also generated. To compute centroid structure we used McCaskill’s algorithm [30] through -p option.

##### Micro-RNA mir-1825 Structural Modelling

We have used SimRNA [31] Tool for simulating circular structure of miRNA mir-1825 as SimRNA generates a circular starting conformation with the 5’ and 3’ ends close to each other as a starting structure for simulation. After specifying Secondary structure restraints using multiline dots-and-brackets format, the dots-and-brackets input is parsed and internally converted into the dedicated list of restraints. W used a default of 500 steps and 1% of the lowest energy frames taken to clustering.

#### RNA Interaction Modeling

IntaRNA [32] is a program for fast and accurate prediction of interactions between two RNA molecules. It has been used to predict mRNA circRNA sites, to represent the interaction energy in the RNA Sponge.

#### Cas9 Modeling

Cas9 of S.pyogenes (BBa_K1218011) part was translated and modelled by SWISS Model server [33, 34] Using SWISS-Model web server the modelling process was initiated by template recognition process where templates were selected according to the maximum sequence similarity 5FQ5 was of highest sequence identity (Sequence identity: 100.00), Finally, the geometry of the resulting model is regularized by using a force field. The global and per-residue model quality has been assessed using the QMEAN scoring function. For improved performance, weights of the individual QMEAN terms have been trained specifically for SWISS-MODEL. Models were selected based on their sequence identity as well as Swiss-MODEL quality assessment parameters GMQE and QMEAN4.

#### Nucleic acid Modelling

The most stable 2D structure of gRNA was generated using vfold [35]. The Rosetta package FARFAR [36, 37] was used to build the 3D structure of gRNA, 3D-DART [38] was used to generate a 3D structure of the target DNA representing the cleavage site and PAM of cas9, while PAM flexibility was studied using Naflex [39].

#### Docking protocol

Following HADDOCK [40, 41] docking protocol, consisting of randomized orientations and rigid body energy minimization, we have calculated 1,000 complex structures. The 200 complexes with the lowest intermolecular energies have been selected for semi-flexible simulated annealing in torsion angle space. The resulting structures have been then refined in explicit water. Finally, the solutions have been clustered using a threshold value of 1.5 Å for the pairwise backbone RMSD at the interface, and the resulting clusters have been ranked according to their average interaction energy (defined as the sum of van der Waals, electrostatic and AIRs energy terms) as well as buried surface area. HADDOCK scoring is performed according to the weighted sum (HADDOCK score) of different energy terms which includes van der Waals energy, electrostatic energy, distance restraints energy, direct RDC restraint energy, intervector projection angle restraints energy, diffusion anisotropy energy, dihedral angle restraints energy, symmetry restraints energy, binding energy, desolvation energy and buried surface area. One lowest energy structure of the lowest intermolecular energy cluster was selected for analysis. This lowest energy structure displayed no AIR restraint violations within 0.3 Å threshold and was accepted as the final docked structure for the complex.

#### Network Modeling

Regulatory interactions between circRNA-miRNA and miRNA-mRNA, were predicted computationally by Targetscan [24] and miRanda algorithm [42], then integrated to represent the regulatory network of ceRNA network of circRNA hsa-circ-0000064 which is HCC related. Both species have been subjected to adjacency matrix set up based on the weight of interaction between predicted targets, circRNA hsa-circ-0000064 and its’ target miRNAs as well as miRNA mir-1825 and its target mRNAs were represented by weight matrix describing relation strength from selected databases. Finally, the final proposed ceRNA network was constructed using Cytoscape 3.5.1 [43]. Network analysis suggested the TRIM2 tumor suppressor which we used to experimentally assess the regulatory function of ceRNA network.

### Wet Lab Methods

We wish to build a single expression plasmid that can express hsa_circ_0000064, we created 2 methods for biobrick assembly to tackle this issue; CRISPR-and non-CRISPR-based synthetic constructs with both genetic constructs consisting of 5 fragments. We retrieved all the required parts from IGEM parts registry, our new parts were submitted to the parts registry and the genetic constructs were designed computationally using benchling framework.

#### Competent E coli Transformation and plasmid DNA purification

We have managed to transform our competent cell colonies with pcDNA™3.1(+) vector containing either CRISPR or direct genetic construct containing hsa_circ_0000064 and miRNA binding site. With these cell lines we will be able to make more copies of each part in preparation for the arrival of our synthesized sequences. Purification of plasmid DNA was done according to Monarch® Plasmid Miniprep Kit (NEB #T1010) Protocol.

#### HepG2 transfection with CRISPR and non-CRISPR-based genetic construct or empty vector

First genetic construct was introduced into cells using LipofectamineTM 2000 reagent (Invitrogen, Carlsbad, CA, USA) according to manufacturer’s instructions. Second CRISPR-based genetic construct was introduced into cells simultaneously with pFETCh_vector expressing hsa_circ_0000064 and HDR using LipofectamineTM 2000 reagent.

#### Viability assay

Cell viability assay was performed by means of a cell counting kit (CCK-8; Dojindo, Kumamoto, Japan). A 96-well plate containing pre-cultured cells (3,000 cells/well) was subjected to media replacement by the WST-8 reagent (2-(2-methoxy-4-nitrophenyl)-3-(4-nitrophenyl) −5–2, 4-disulphonyl)-2H-tetrazolium monosodium salt) (which when reduced turns to orange formazan) at the indicated time points. Both the developed color and the absorbance were measured at 450 nm, with the amount of formazan being directly proportional to the number of living cells, using a microplate reader (NEC, Tokyo, Japan).

#### QRT-PCR for circular RNA –miRNA –mRNA genetic network in human sera samples

RT2 miRNA First Strand Kit (Qiagen, Valencia, CA) using miScriptHiSpec buffer was used to prepare cDNA from 1 µg RNA. Real-time PCR was performed using a Step one Plus Applied Biosystem. All reactions were performed in triplicates.

hsa-circ-0000064 and TRIM mRNA expression in HCC cell lines were assessed using QuantiTect SYBR Green PCR Kit (Qiagen, Valencia, CA) and gene specific primers (Circular RNA specific Quantict Primer and Hs_TRIM2_1_SG Quantict Primer), on Step One Plus™ System (Applied Biosystems Inc., Foster, CA). Beta actin was used as a housekeeping gene. Divergent primers forhsa_0000064 primers were designed by Circinteractome datatabase (“http://circinteractome.nia.nih”) and synthesized by (Qiagen, Germany).

MiR-1825 expression in HCC cell line was assessed by mixing the total cDNAs with miRNA-specific forward primer (Hs_miR-1825_1 miScript Primer Assay miScript Primer Assay) and miScript SYBR Green PCR Kit (Qiagen/SABiosciences Corporation, Frederick, MD) according to the manufacturer’s protocol. RNU-6 was used as an internal control

The PCR program for Syber green based QPCR was as follow: denaturation at 95°C for 15 min; followed by 40 cycles of denaturation for 10 sec at 94°C; then annealing for 30 sec at 55°C; finally, extension for 34 sec at 70°C. Each reaction was done in duplicate. Relative quantification of gene expression was calculated using the 2-ΔΔCt method. The cycle threshold (Ct) value of each sample was calculated using Step One Plus™ software v2.2.2 (Applied Biosystems). Any Ct value above 36 was considered negative. Amplification plots and Tm values were performed to ensure the specificities of the amplicons.

#### Flow Cytometery and Immunocytochemical staining (ICC)

Single cell suspension was prepared from harvested cultured cells, fixed with100ul immunofixation buffer, then incubated at room temperature for 20-30 minutes protected from light after permeabilization by permibilization buffer, cells were washed with PBS and labelled with TRIM2 antibody (PA5-57431). Goat anti-Rabbit IgG (H+L) Highly Cross-Adsorbed Secondary Antibody Alexa Fluor® 488 conjugate (A-11034) was used at a concentration of 4 μg/ml and analysed by FlowCytometryFor immunostaing, adherent HepG2 cells were fised then stained as above. No nonspecific staining was observed with the secondary antibody alone, or with an isotype control. The images were captured at 60X magnification. ICC staining of HepG2 cells after co-expression of hsa-circ-0000064 expressing vector shows the brightness of TRIM2 protein prominently in the HepG2 cells transfected with either CRISPR-or non-CRISPR-based genetic construct.

## Results

### Dry Lab Results

#### Biomarker Filtration Results

Our modeled ceRNA network contained, the circular RNA (has-cric-0000064) competing for shared microRNA (mir-1825) and sequestrate it within the cell as they have MREs (microRNA sponge); lastly deregulating our target gene (TRIM2).

#### Assembly of genetic constructs

Our design aims to compare the function of both circuits at regulating non-coding RNAs

The first CRISPR circuit submitted as BBa_K2217026 part and ceRNA circuit submitted as BBa_K2217025 part.

**Figure-1.**
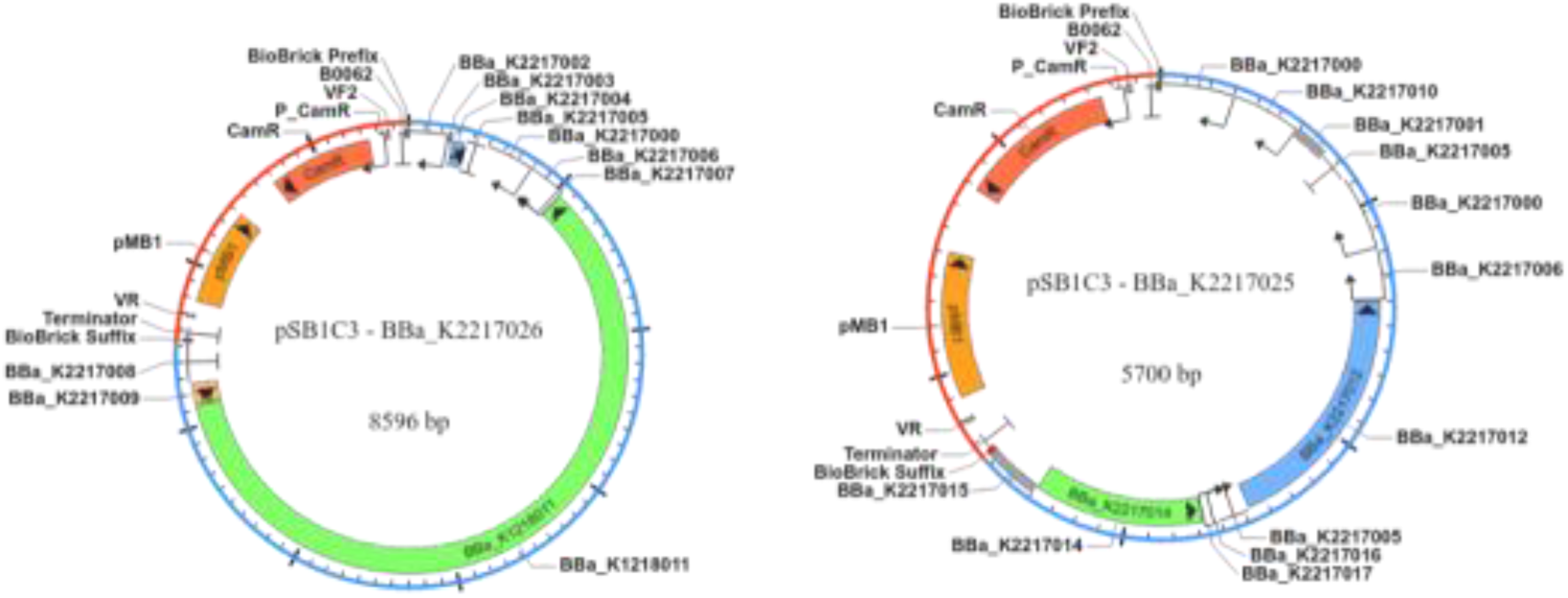
shows our two main circuits submitted as separate parts the first CRISPR circuit submitted as BBa_K2217026 part and ceRNA circuit submitted as BBa_K2217025 part.

For constructing CRISPR-based circuit submitted as BBa_K2217026 part, Part BBa_K2217018 was composed of U6 (BBa_K2217002) as a promoter for gRNA (BBa_K2217003) (Designed using MIT CRISPR web tool [44] The gRNA needs a scaffold to guide the Cas9 (BBa_K2217004) and finally it was terminated by SV40 poly(A) signal Termination (BBa_K2217005). Part BBa_K2217019 was composed of CMV Enhancer as well as CMV Promoter (BBa_K2217000 and BBa_K2217006 respectively) as constitutive promoters to enhance the transcription process of the Cas9 (BBa_K1218011) part which was transcribed using T7 Promoter (BBa_K2217007). The design aimed at generating Homology Directed Repair instead of Non-Homologous end joining repair of Cas9 by knocking in the circular RNA as a Competing endogenous RNA enhancing its transcription to regulate miRNA action, so we designed Homology repair template (BBa_K2217009) to be transfected on a separate donor vector then terminated with CYC1 terminator (BBa_K2217008).

In order to construct ceRNA-based circuit submitted as BBa_K2217025 part, we used the CAG promoter composed of CMV enhancer and Chicken B-actin Promoter (BBa_K2217013) while circular RNA (hsa-circ-0000064) was submitted to the registry after biomarker filtration using bioinformatics databases as (BBa_K2217001). Finally it was terminated by SV40 poly (A) signal Termination (BBa_K2217005). Part BBa_K2217022 was composed of CMV Enhancer as well as CMV Promoter (BBa_K2217000 and BBa_K2217006 respectively) as constitutive promoters to enhance the transcription process of the laci that was submitted (Ba_K2217012). We also improved the fragment’s characterization by adding the miRNA binding site of the miRNA mir-1825 as determined computationally using circInteractome database [23] where we hypothesized its function as a trigger for miRNA binding and synthesis. Finally, it was terminated by SV40 poly (A) signal Termination (BBa_K2217005). Part BBa_K2217023 is composed of lac operator (BBa_K2217016) and the lac promoter (BBa_K2217017) as a weak constitutive promoter in a trial to improve characterization of ceRNA network using yfb (BBa_K2217014) detection using flow Cytometery. Finally, it was terminated by Poly_gh termination (BBa_K2217015).

### Modeling Results

#### CeRNA network Deterministic Model results

Our Model aims to describe the regulation of competing endogenous RNA (ceRNA) network using ordinary differential equations to get insights about the kinetics of molecular species inside the network. The Model was constructed in Synthetic biology markup language SBML. SBML models were converted to SBOL (synthetic biology open language) to describe biological parts and their interactions including: transcription, degradation, association and dissociation of both the ceRNA and miRNA. The Model describes an inhibitory relationship, where the miRNA binding to ceRNA inhibits the miRNA action on its target mRNA. We can estimate that effect from the change in free miRNAs in the simulation run. Parameters have been estimated from the work of Bosia et al. [27] and described as a system of ODEs.

**Figure-2.**
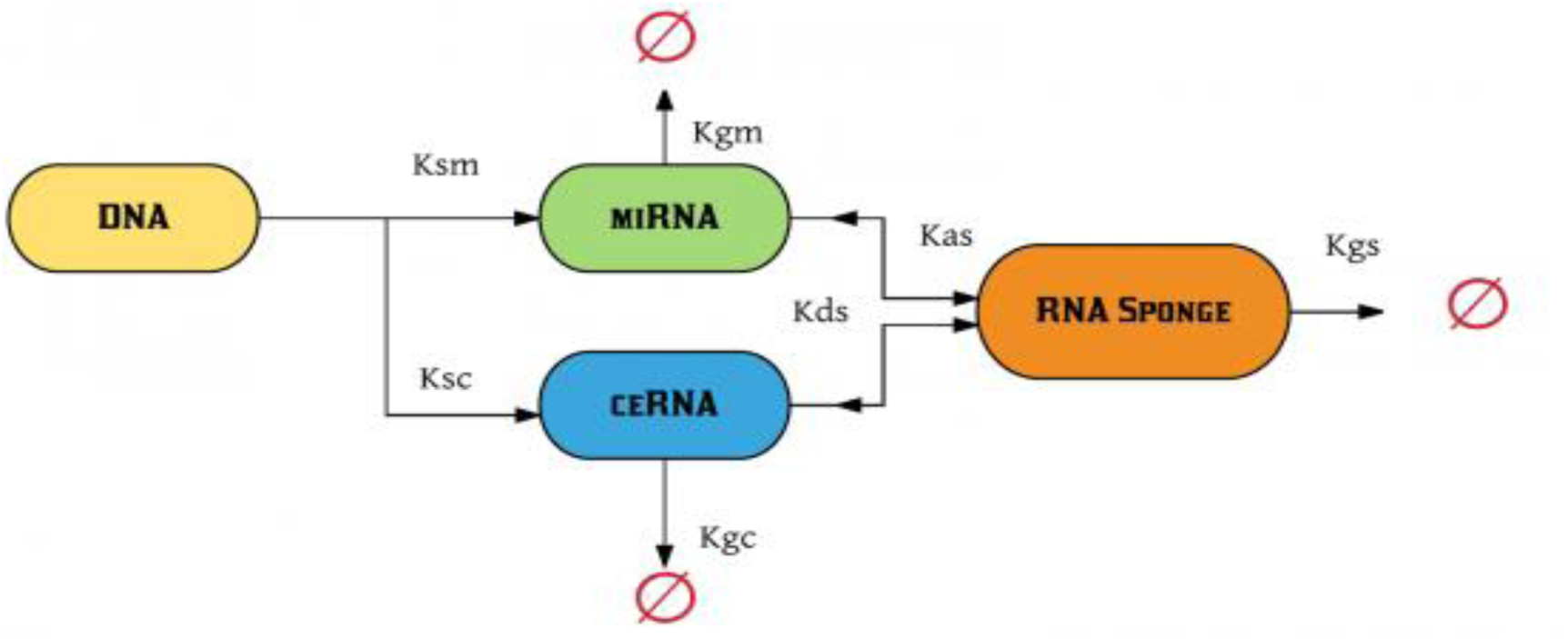
Graphical representation for ceRNA model

**Figure-3.**
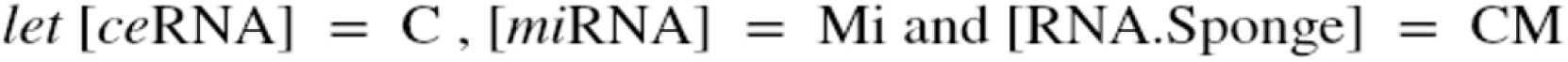
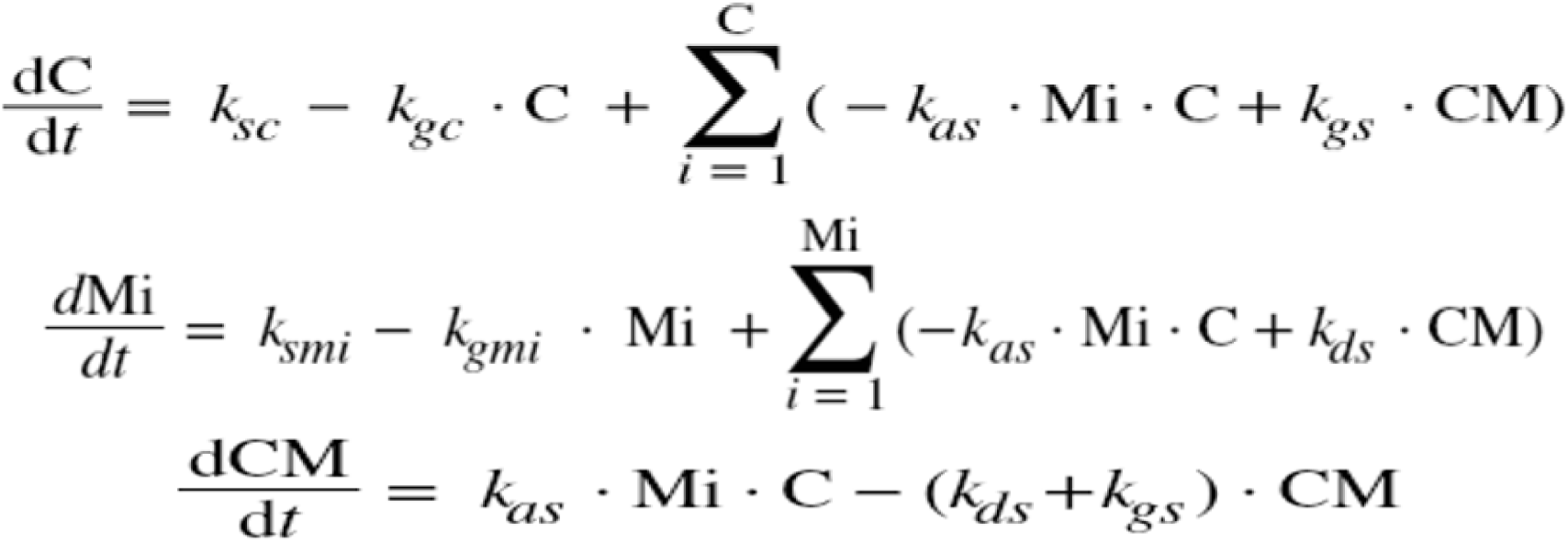
Set of Ordinary Differential Equations ODEs Representing ceRNA network deterministic model according to Bosia et al.

**Table-1.** Parameter values of ceRNA network regulation

| Symbol | Description | Value |
| --- | --- | --- |
| Ksmi | Rate of transcription of miRNA | 0.2s<sup>-1</sup> |
| Ksc | Rate of transcription of ceRNA | 0.155s<sup>-1</sup> |
| Kgmi | Rate of degradation of miRNA | 0.0003s<sup>-1</sup> |
| Kgc | Rate of degradation of ceRNA | 0.0004s<sup>-1</sup> |
| Kas | Rate of association of RNA Sponge Complex | 0.0005s<sup>-1</sup> |
| Kds | Rate of dissociation of RNA Sponge Complex | 0.0003s<sup>-1</sup> |
| Kgs | Rate of degradation of RNA Sponge Complex | 0.00031s<sup>-1</sup> |

**Figure-4.**
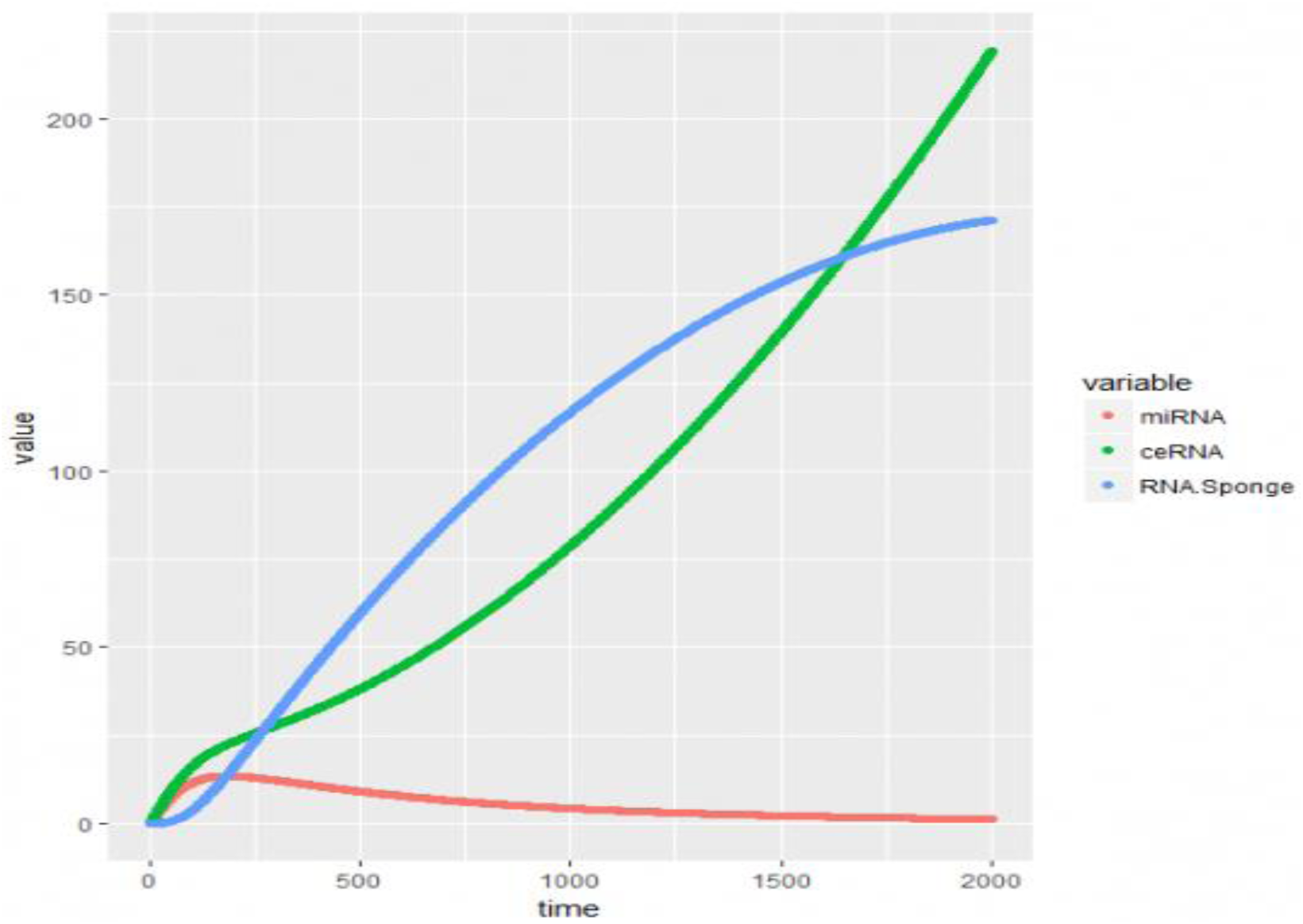
Simulation run for ceRNA network model ODEs

The simulation in Figure-4 Shows inhibitory relationship along time axis to the quantity of free miRNA along the transcription of circular RNA as a competing endogenous RNA which may describe the sponge action, regarding the elevation of free miRNA action on target mRNAs.

#### CRISPR Deterministic Modeling Results

Our Model aims to describe the regulation of CRISPR network. The Model was constructed in Synthetic biology markup language SBML, including: transcription, degradation, and association of gRNA and cas9. The model describes the binding interaction between the gRNA and cas9 that is supposed to be informative to the cas9 about the cleavage site near the PAM. Parameters have been estimated from the work of R.moore et al [46].and described as a system of ODEs.

**Figure-5.**
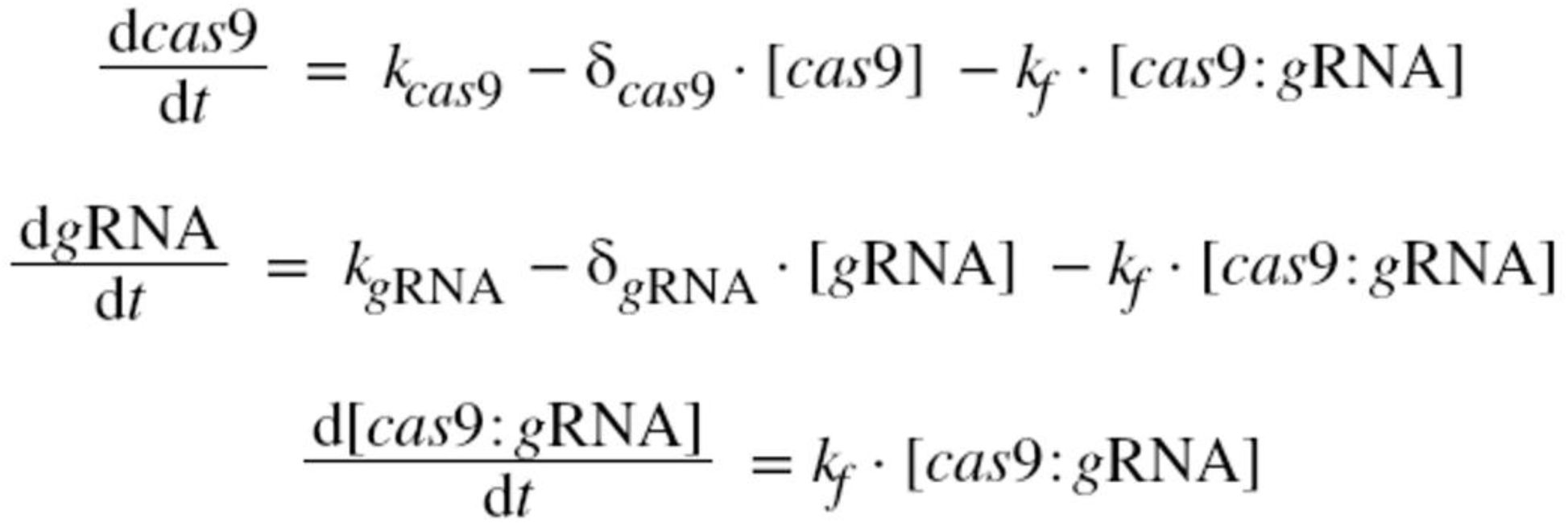
Set of Ordinary Differential Equations ODEs Representing CRISPR network deterministic model

**Figure-6.**
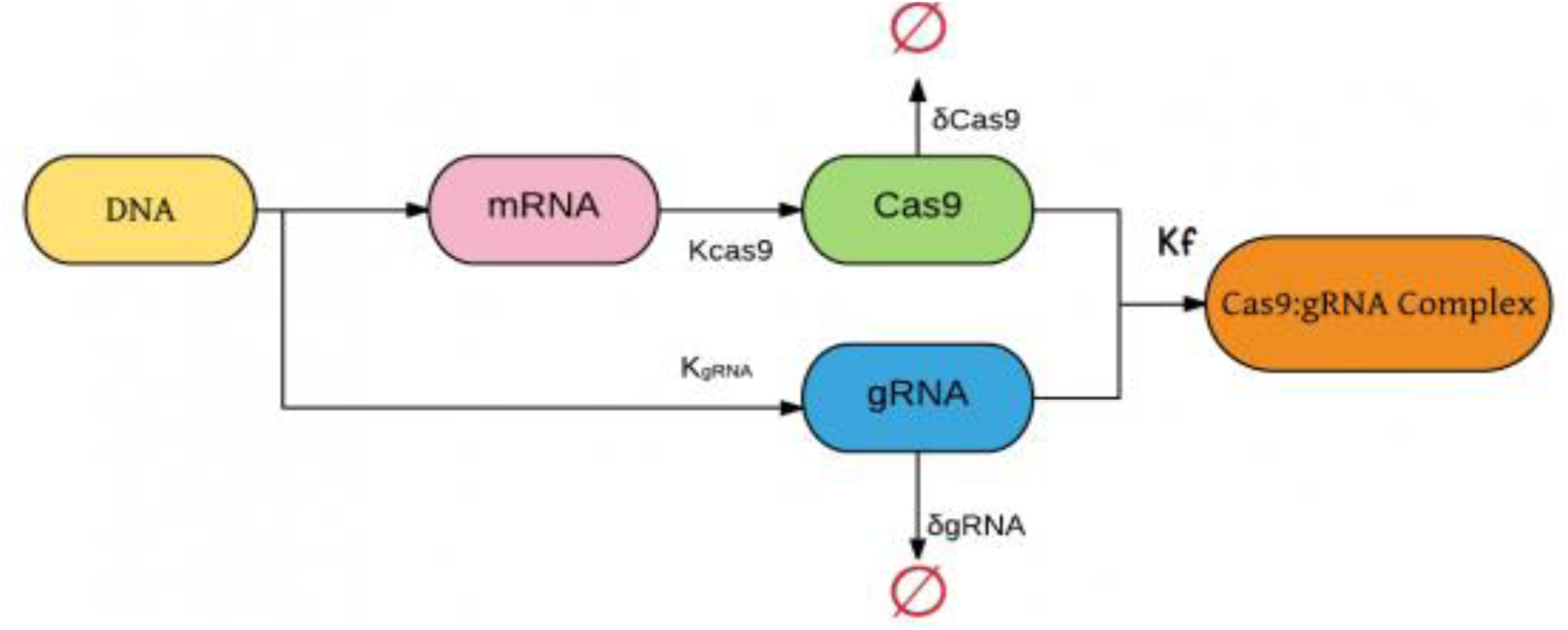
Graphical representation for CRISPR network deterministic model (Moore et al.)

**Table-2.**
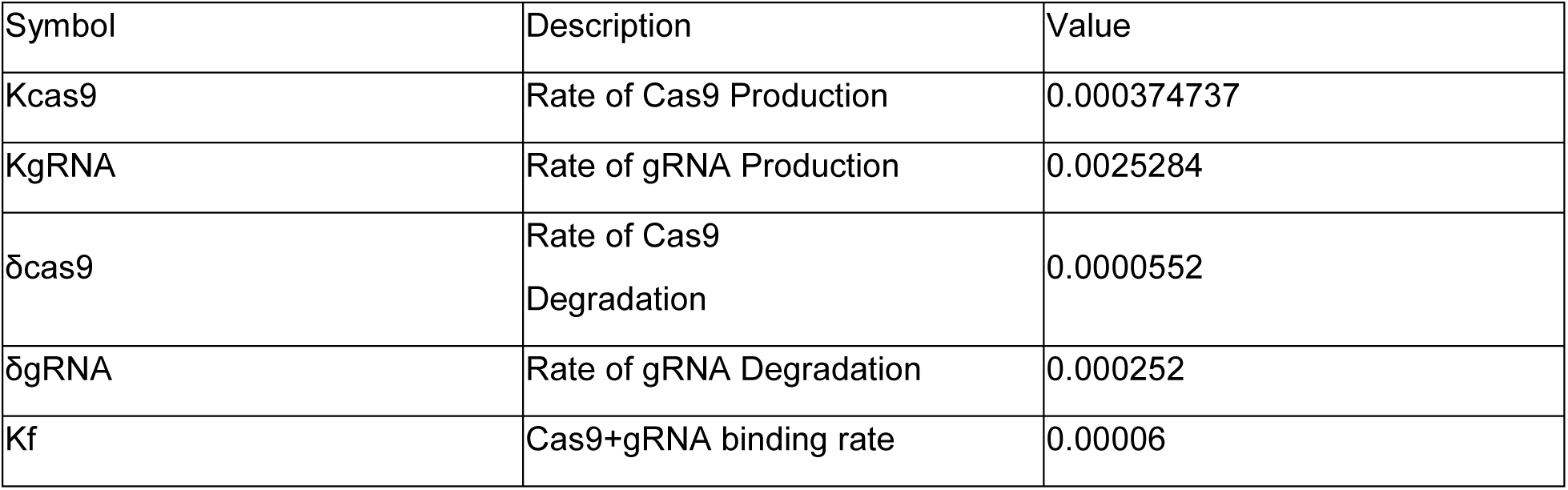
Parameter values of cas9 network regulation

| Symbol | Description | Value |
| --- | --- | --- |
| $K_{cas9}$ | Rate of Cas9 Production | 0.000374737 |
| $K_{gRNA}$ | Rate of gRNA Production | 0.0025284 |
| $\delta_{cas9}$ | Rate of Cas9 Degradation | 0.0000552 |
| $\delta_{gRNA}$ | Rate of gRNA Degradation | 0.000252 |
| $K_f$ | Cas9+gRNA binding rate | 0.00006 |

**Figure-7.**
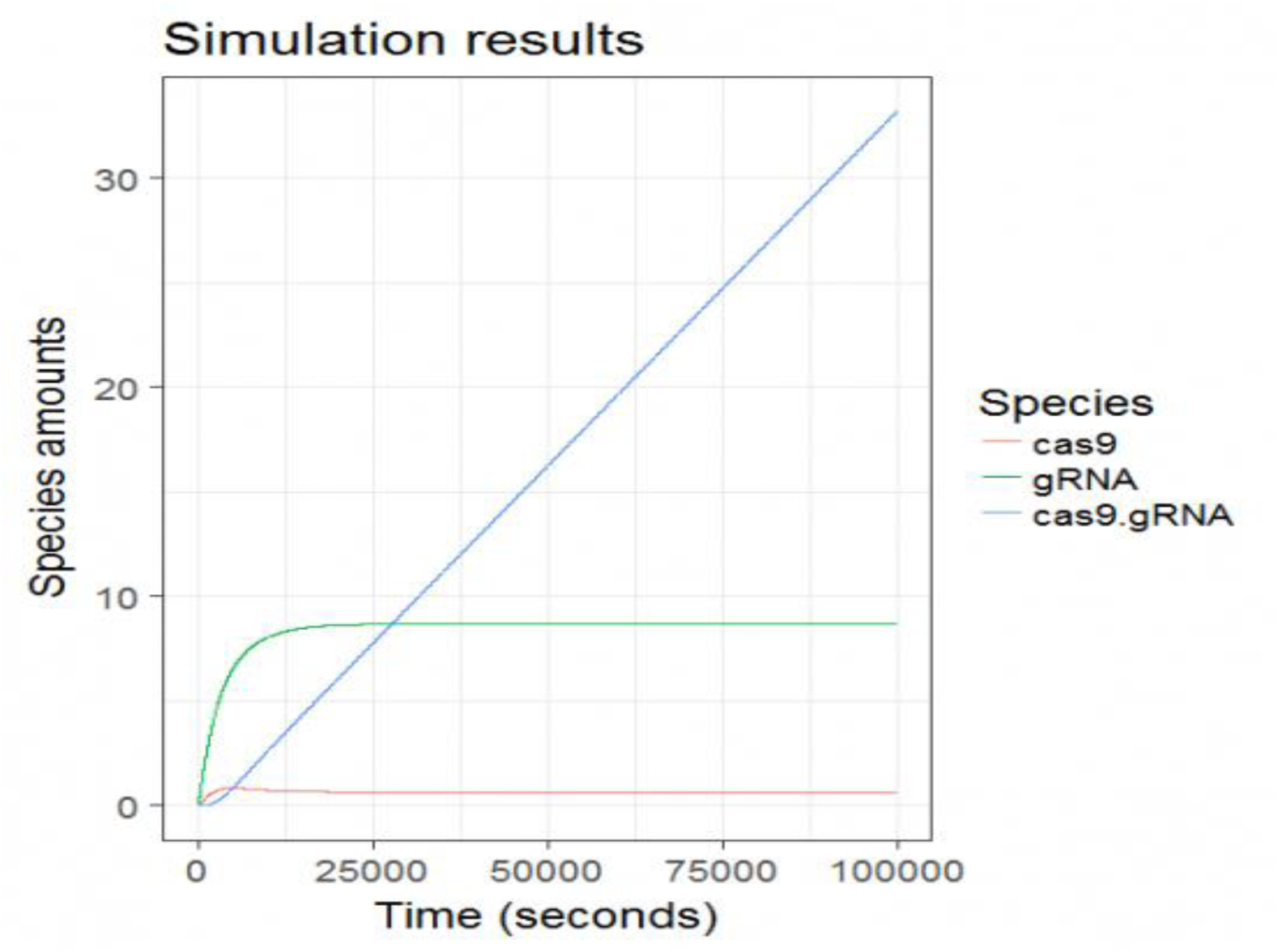
Simulation run for CRISPR network model

The simulation in Figure-7 describes interaction association between gRNA and cas9 along time axis to the quantity of formed complexes cas9. gRNA along the transcription of gRNA as a directing RNA molecule which may describe the complex action regarding focusing cas9 cleavage action on target DNA.

#### Stochastic Modelling Results

Simulations were plotted (Figure-8, Figure-9) to visualize species evolution overtime. These plots were used to assess the accuracy of stochastic modelling using ssa function by comparing stochastic models to deterministic modeling simulations. We have used the function solveStoch of sysBio R package which allowed us to simulate the model using Gillespie stochastic simulation algorithm through the “ssa” function of the GillespieSSA package. A vector was created for modeling parameters. Species, rates and parameters of each model were checked. The list of reactions for stochastic simulation was prepared as a propensity function list. Stochastic matrix was generated where each column represents a reaction while rows represents reaction species. A vector of propensity functions was created. Finally, models were solved stochastically.

**Figure-8.**
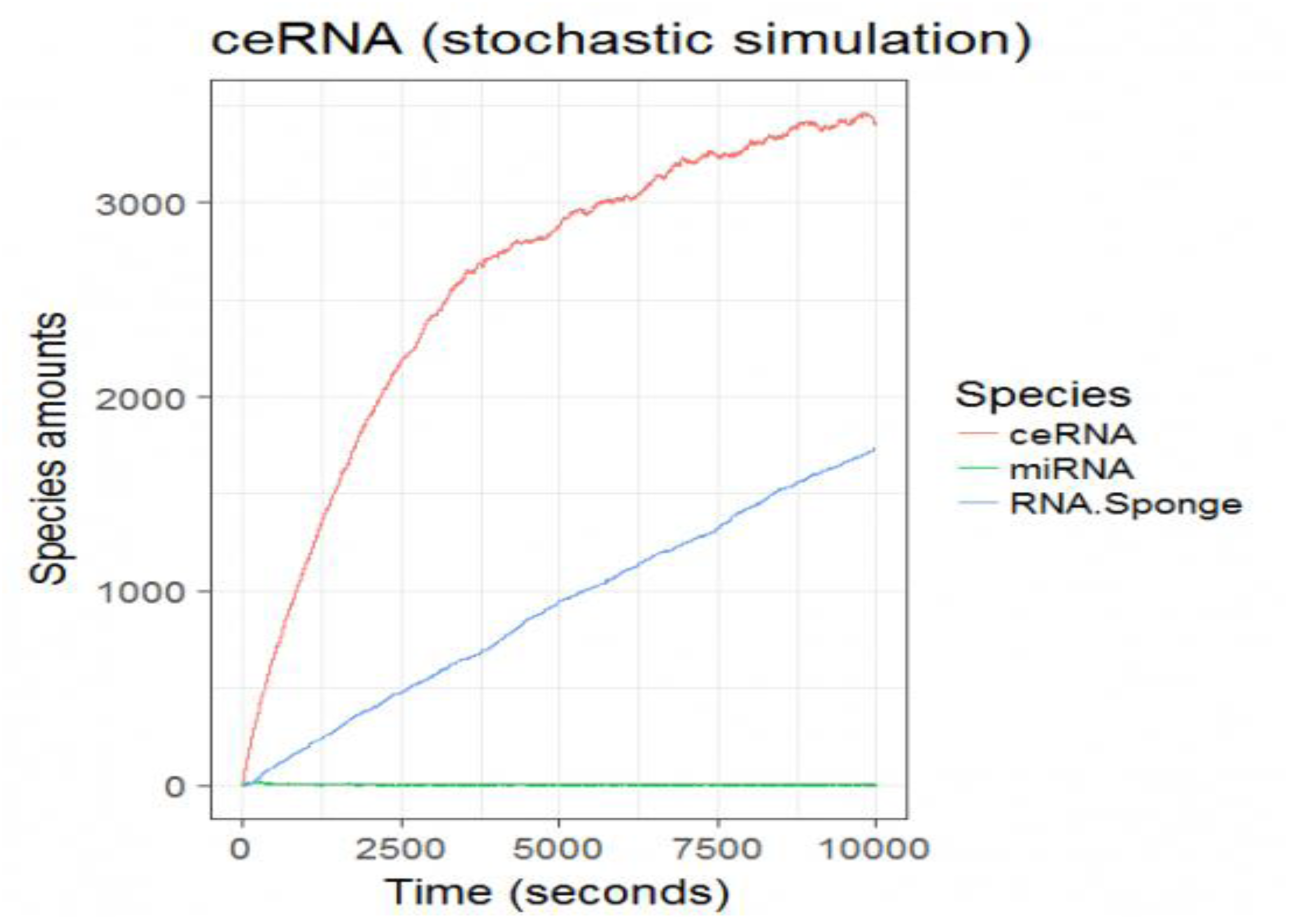
Stochastic Simulation run for ceRNA network model using GillespieSSA [45] R package. This simulation was run using sysBio [25] R package describing stochastic effects on ceRNA network Sponge production and species levels in the network.

**Figure-9.**
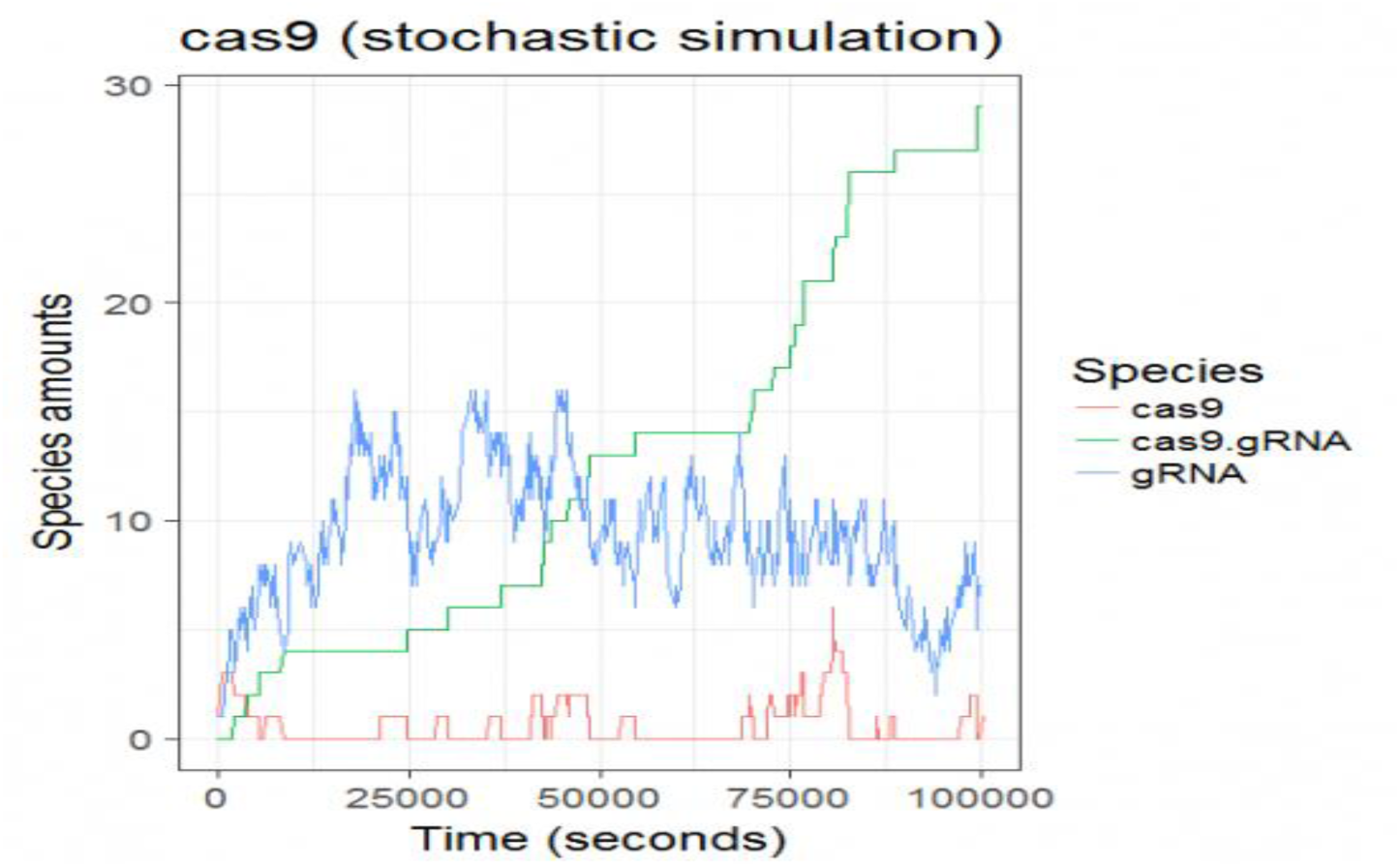
Stochastic Simulation run for CRISPR network model using GillespieSSA [45] R package. This simulation was run using sysBio [25] R package describing stochastic effects of gRNA.cas9 association to form their intermediate complex.

#### Structural Modeling Results

**Figure-10.**
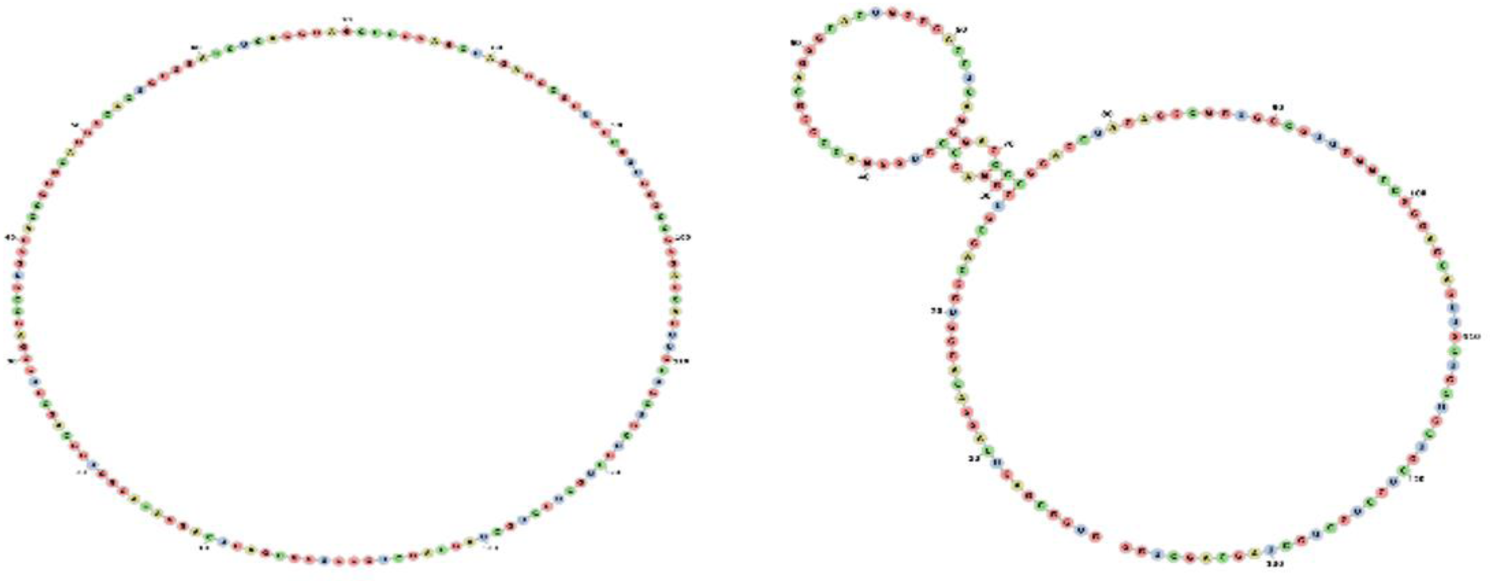
RNAFOLD simulation of circular RNA structure

Minimum free energy prediction using RNAFOLD generated an optimal secondary structure in dot-bracket notation from a centroid structure of 0.00 kcal/mol minimum free energy to 1.78 kcal/mol. Thermodynamic ensemble prediction using RNAFOLD computed a free energy of −51.72 kcal/mol, The frequency of the MFE structure in the ensemble is 0.07 % and the ensemble diversity is 65.27.

**Figure-11.**
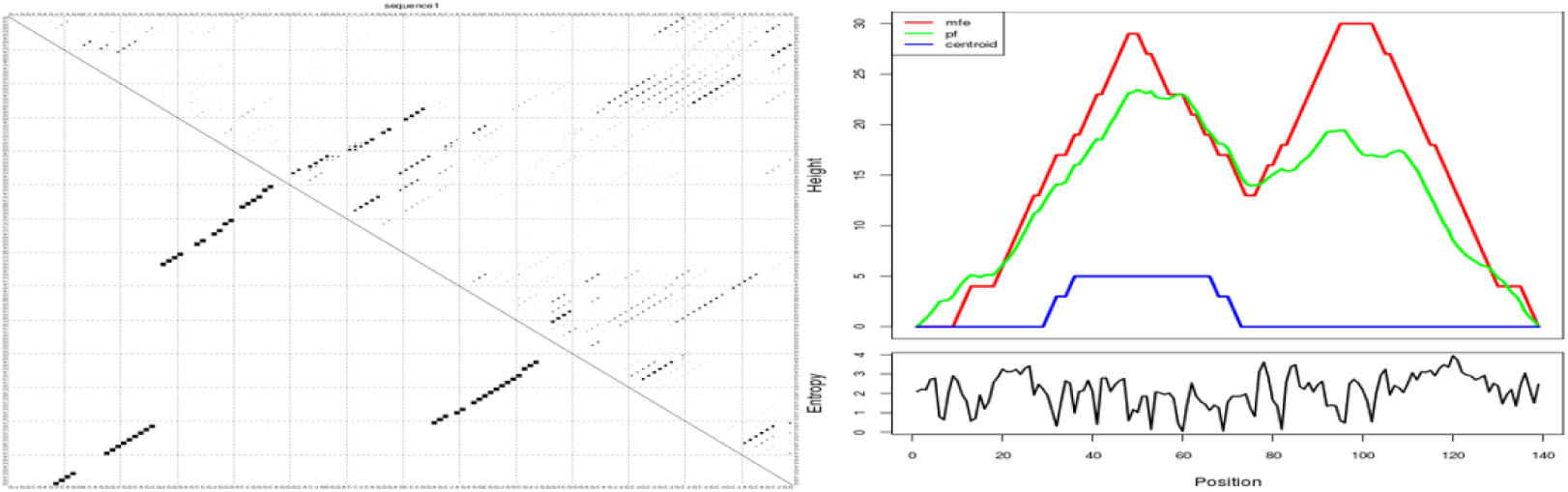
(left) energy dot-plot of circRNA model, (Right) Mountain plot of the same model

To the left is the energy dot 2D plot which indicates all of the base pairs involved in optimal and suboptimal secondary structures, both axes of the graph represent the same RNA sequence. Each point drawn indicates a base pair between the ribonucleotides whose positions in the sequence are the coordinates of that point on the graph. To the right is the mountain plot plotting the number of base pairs enclosing a sequence position versus number of base pairs enclosed at this position. The plot includes the MFE structure (Red), the thermodynamic ensemble of RNA structure (Green), and the centroid structure (Blue). Additionally we used it to estimate the positional entropy for each position.

**Figure-12.**
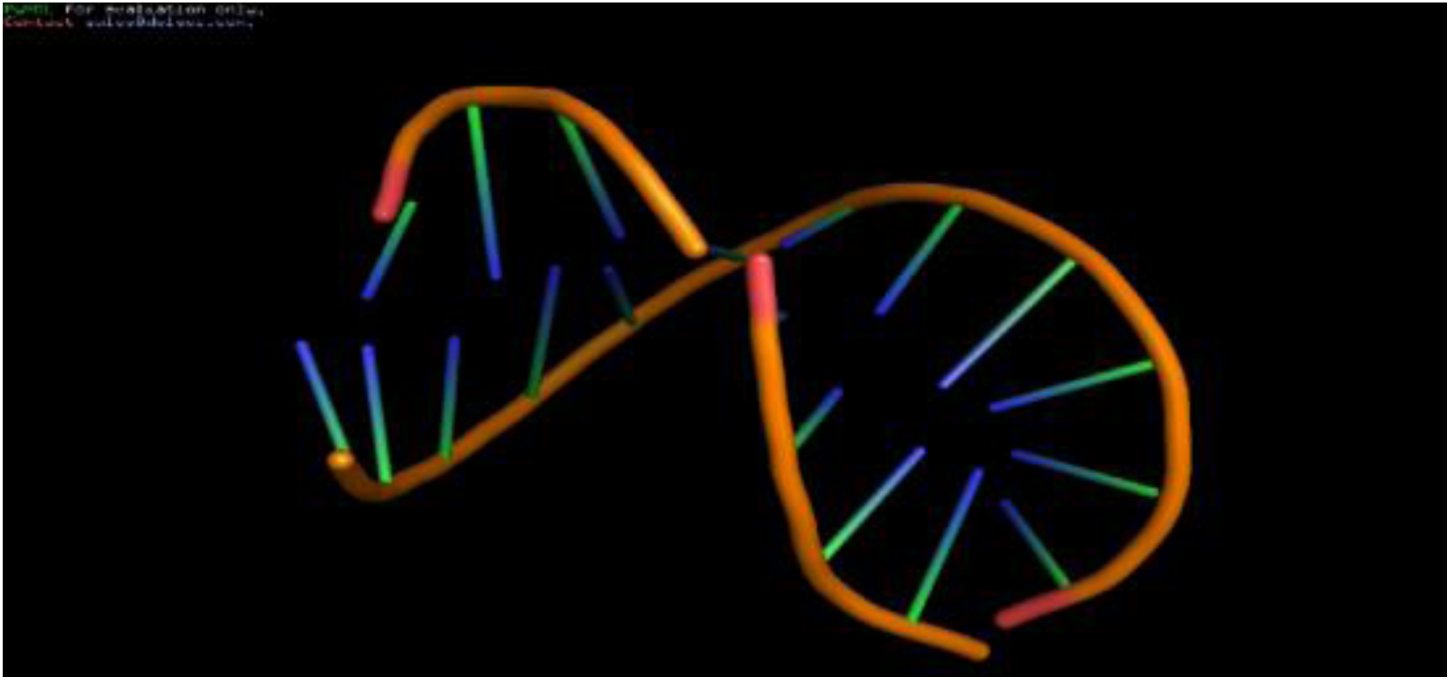
Best Secondary Structure Cluster Predicted by SimRNA for miRNA mir-1825, the back bone colored orange while the riboneucleotides colored green and blue with red capping of the double stranded endings using PyMol visualization software.

**Table-3.**
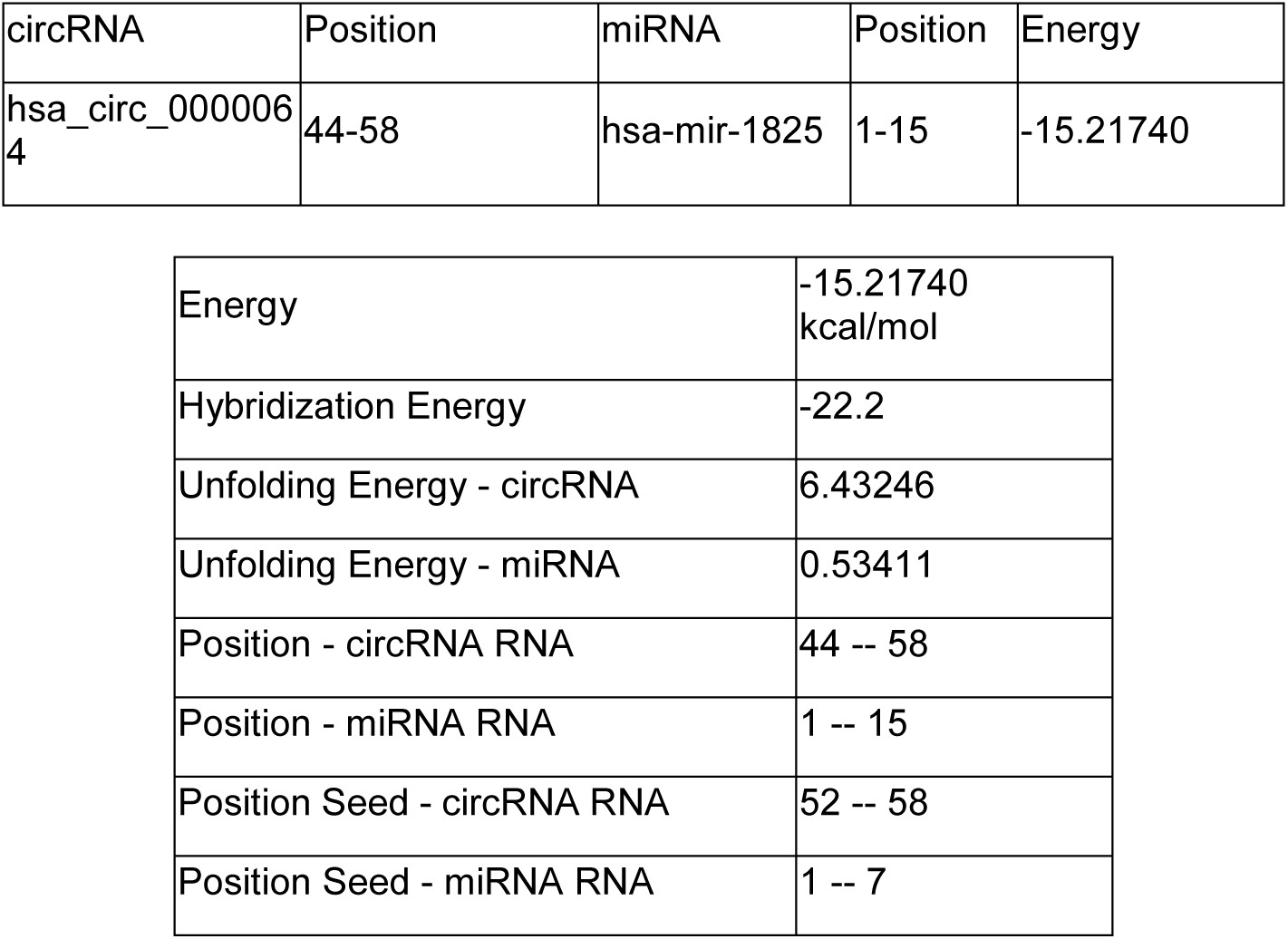
IntaRNA interaction energy prediction of RNA sponge

| circRNA | Position | miRNA | Position | Energy |
| --- | --- | --- | --- | --- |
| hsa_circ_0000064 | 44-58 | hsa-mir-1825 | 1-15 | -15.21740 |

|  |  |
| --- | --- |
| Energy | -15.21740 kcal/mol |
| Hybridization Energy | -22.2 |
| Unfolding Energy - circRNA | 6.43246 |
| Unfolding Energy - miRNA | 0.53411 |
| Position - circRNA RNA | 44 -- 58 |
| Position - miRNA RNA | 1 -- 15 |
| Position Seed - circRNA RNA | 52 -- 58 |
| Position Seed - miRNA RNA | 1 -- 7 |

**Figure-13.**
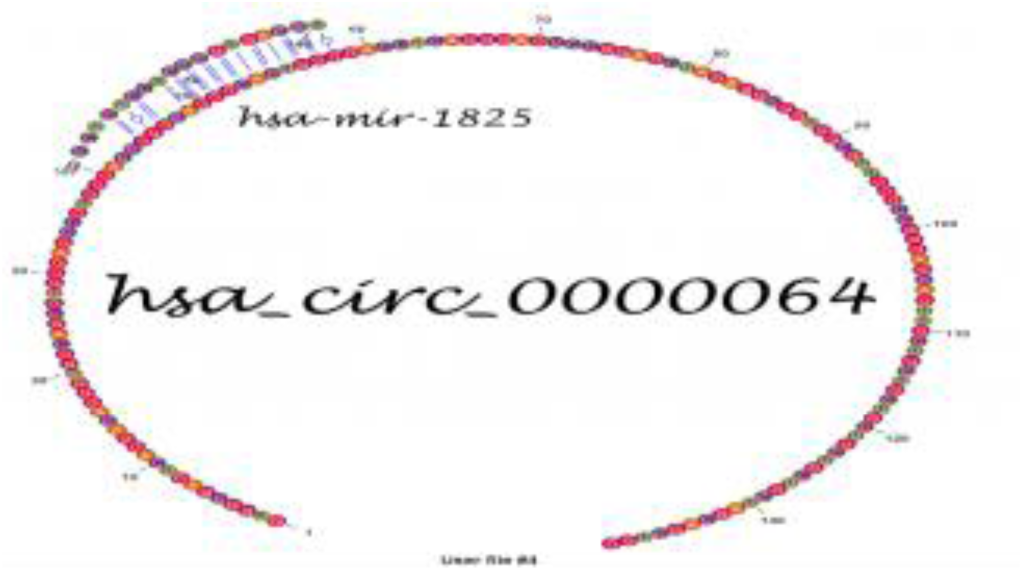
RNA sponge graphical representation of circRNA hsa_circ_000004 and miRNA hsa-mir-1825 in vienna format

#### Position-wise minimal energy profile

The following plots give us insights into the overall circRNA-miRNA interaction abundance. A minimal energy profile is provided for both sequences of both miRNA and circRNA, taking RNA-RNA interaction in consideration.

**Figure-14.**
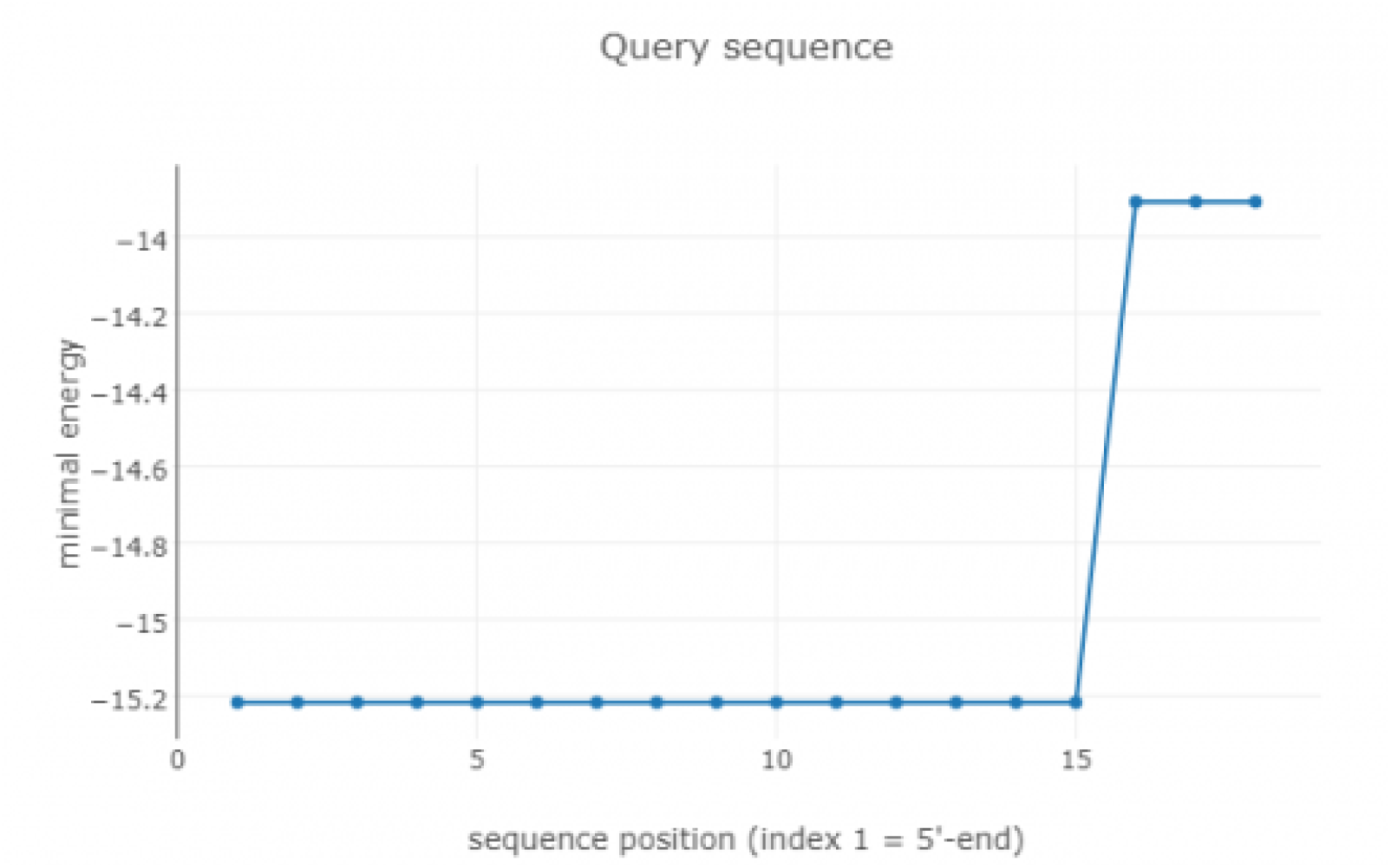
Minimal energy profile is provided for miRNA sequence

**Figure-15.**
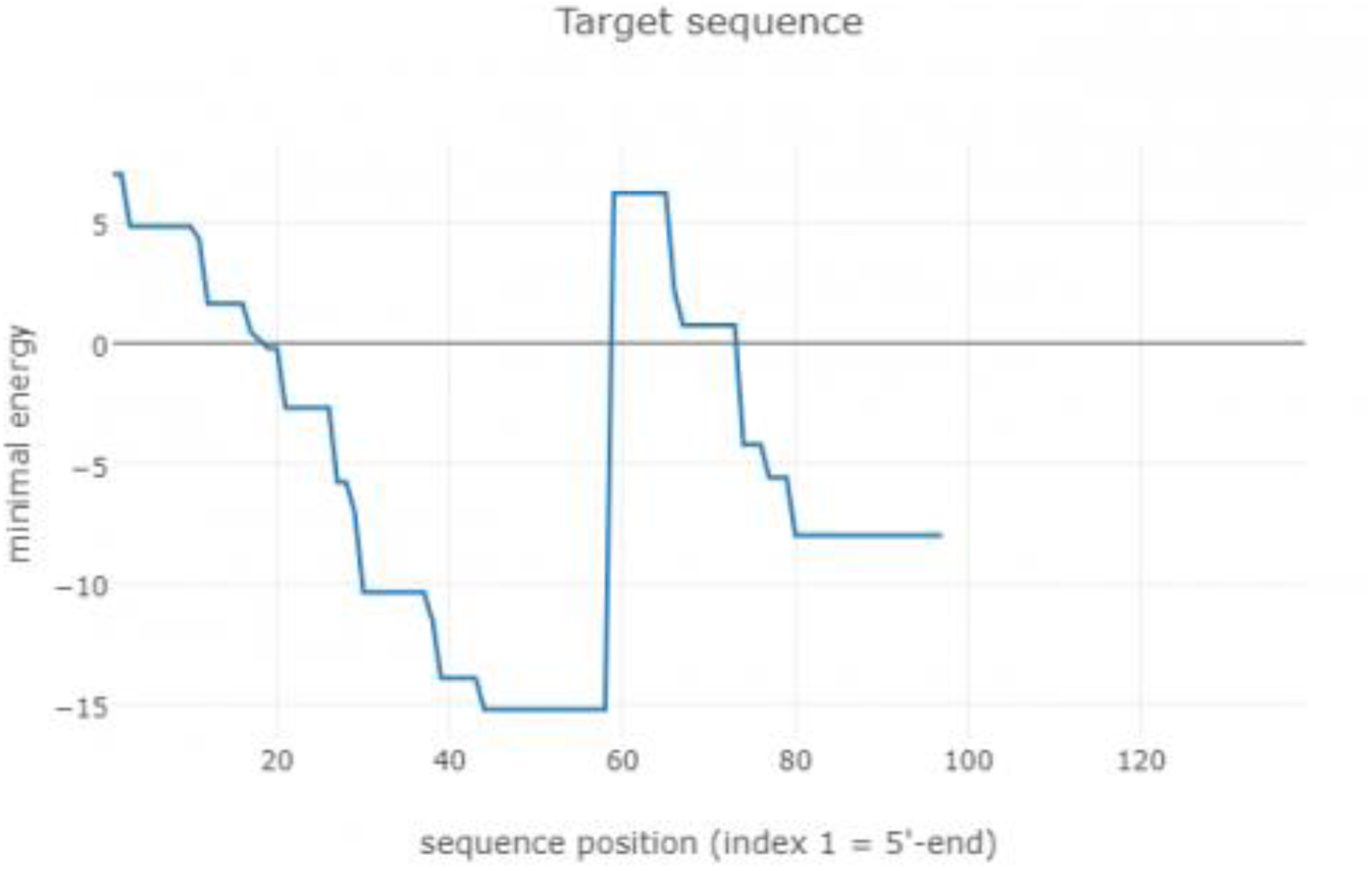
Minimal energy profile is provided for Circular RNA sequence representing sponge interaction.

**Figure-16.**
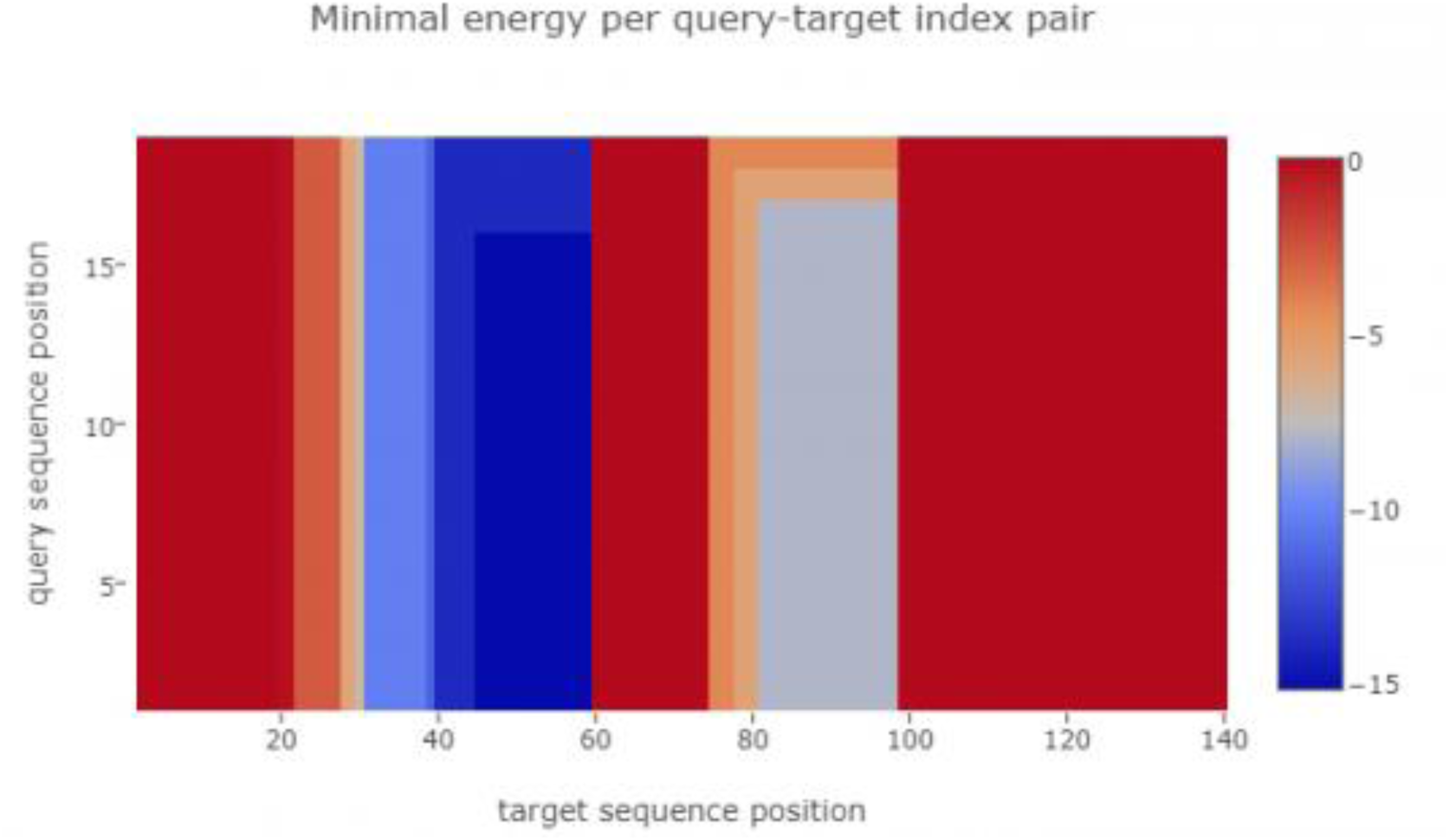
Heatmap visualization of the minimal energy for each non-coding RNA in the sponge

#### Cas9 Modelling results

The geometry of the resulting model is regularized by using a force field. The global and per-residue model quality has been assessed using the QMEAN scoring function. For improved performance, weights of the individual QMEAN terms have been trained specifically for SWISS-MODEL. Models were selected based on their sequence identity as well as Swiss-MODEL quality assessment parameters GMQE and QMEAN4.

**Figure-17.**
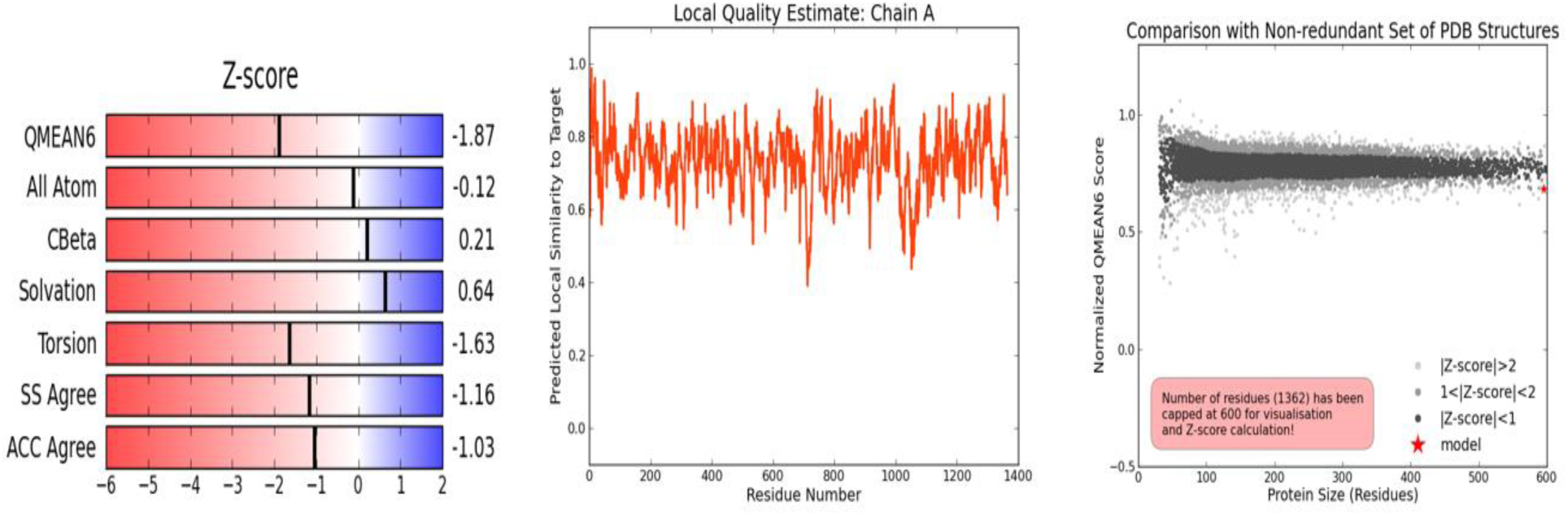
Quality estimation plots based on SWISS-MODEL parameters for Cas9

**Table-4.** Docking scores of both complexes using HADDOCK

| Complex | Cas9.gRNA Cluster | Cas9.gRNA:Target DNA Cluster |
| --- | --- | --- |
| HADDOCK score | -75.64 +/- 22.9 | -340.7 +/- 23.9 |
| clusters | 13 | 17 |
| Total Interaction energy(Kcal mol<sup>-1</sup>) | 1.0 +/- 0.6 | 6.3 +/- 2.1 |
| Van der Waals energy(Kcal mol<sup>-1</sup>) | -66.7 +/- 19.5 | -97.3 +/- 14.1 |
| Electrostatic Energy(Kcal mol<sup>-1</sup>) | -284.4 +/- 21.7 | -342.0 +/- 73.6 |
| Desolvation energy(Kcal mol<sup>-1</sup>) | 20.6 +/- 4.0 | -201.2 +/- 14.6 |
| Restraints violation energy(Kcal mol<sup>-1</sup>) | 273.9 +/- 32.63 | 262.1 +/- 60.51 |
| Buried Surface area | 1912.6 +/- 303.2 | 2785.3 +/- 168.1 |
| z-score | -2 | -2 |

For Cas9.gRNA docking, HADDOCK clustered 105 structures in 13 cluster(s), which represents 52.5 % of the water-refined models that were generated by HADDOCK. Note that currently the maximum number of models considered for clustering is 200. While for Cas9.gRNA: Target DNA docking, HADDOCK clustered 107 structures in 12 cluster(s), which represents 53.5 % of the water-refined models were generated by HADDOCK.

#### Docking Results analysis

**Figure-18.**
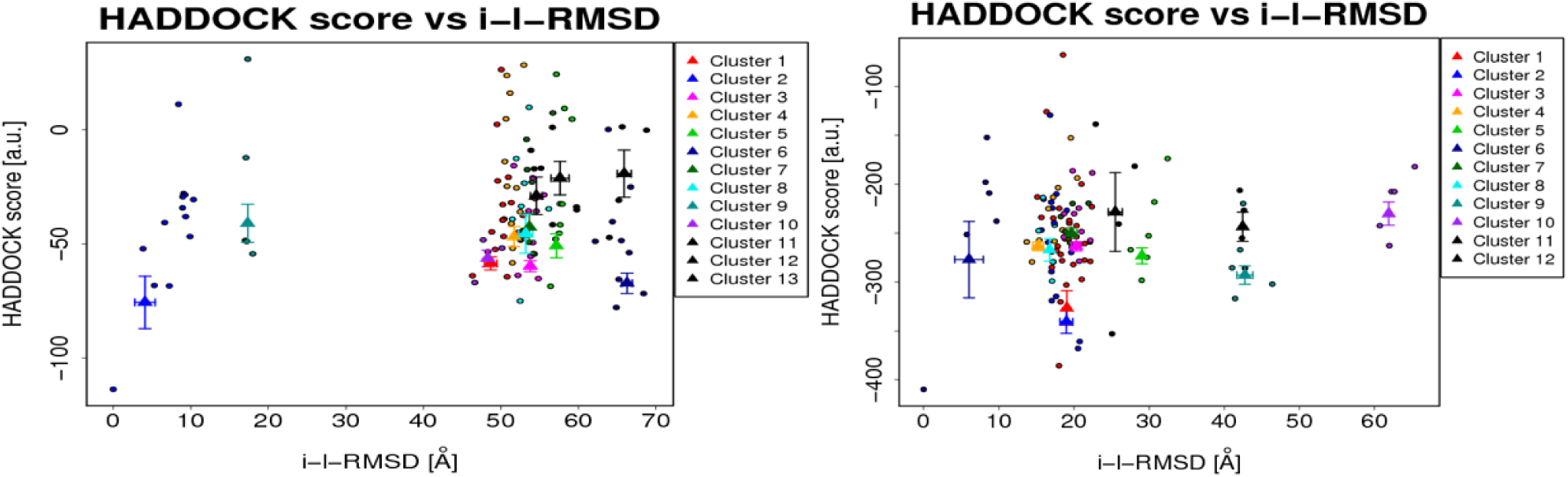
The clusters (indicated in color in the graphs) are calculated based on the interface-ligand RMSDs calculated by HADDOCK, with the interface defined automatically based on all observed contacts to the left is Cas9.gRNA Cluster and to the right is Cas9.gRNA:Target Cluster of lowest energy

**Figure-19.**
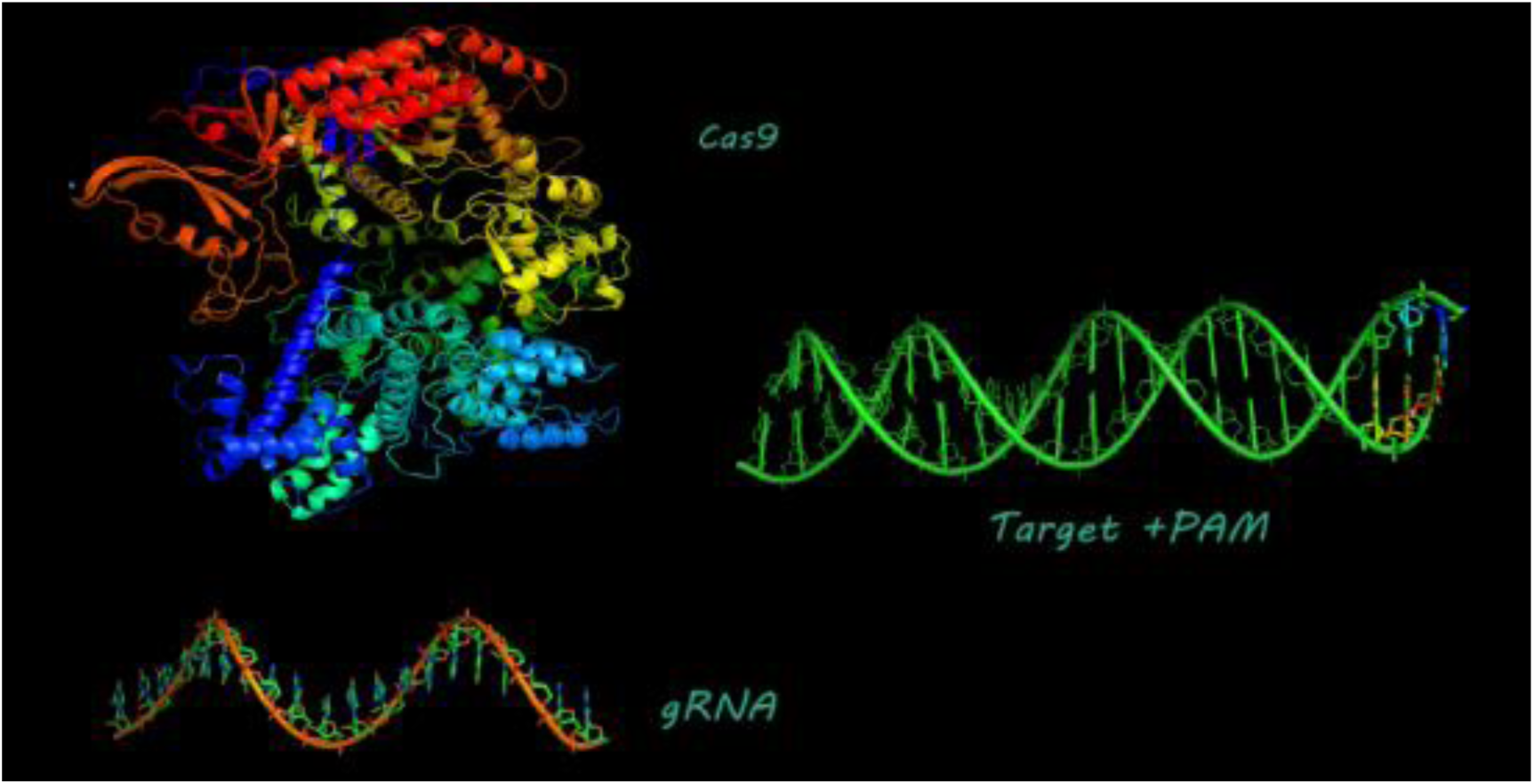
PyMol visualization of Modelled Structures of cas9, gRNA and Target DNA+PAM

**Figure-20.**
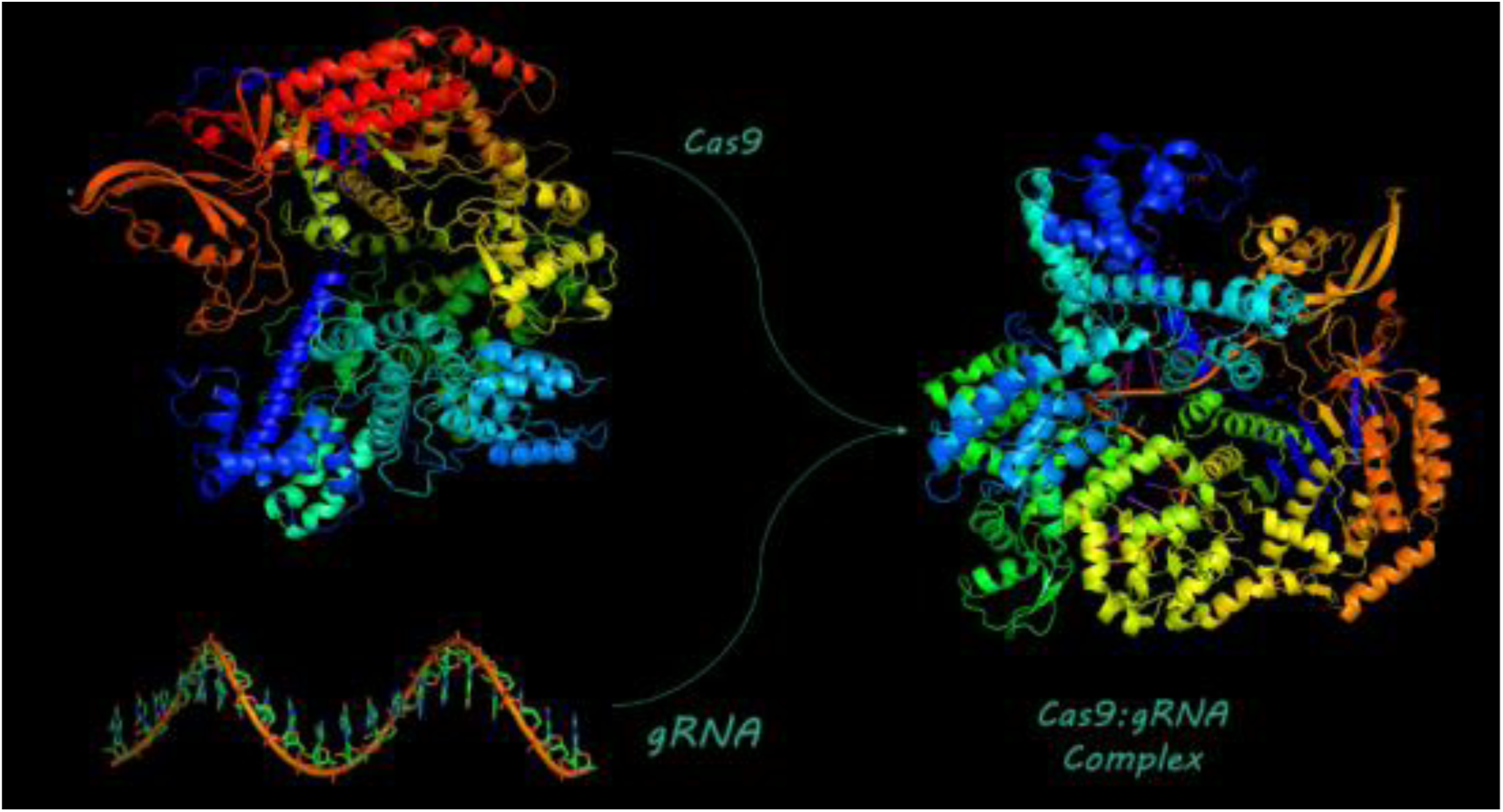
PyMol visualization of Modelled Structures of cas9, gRNA and cas9.gRNA docked complex

**Figure-21.**
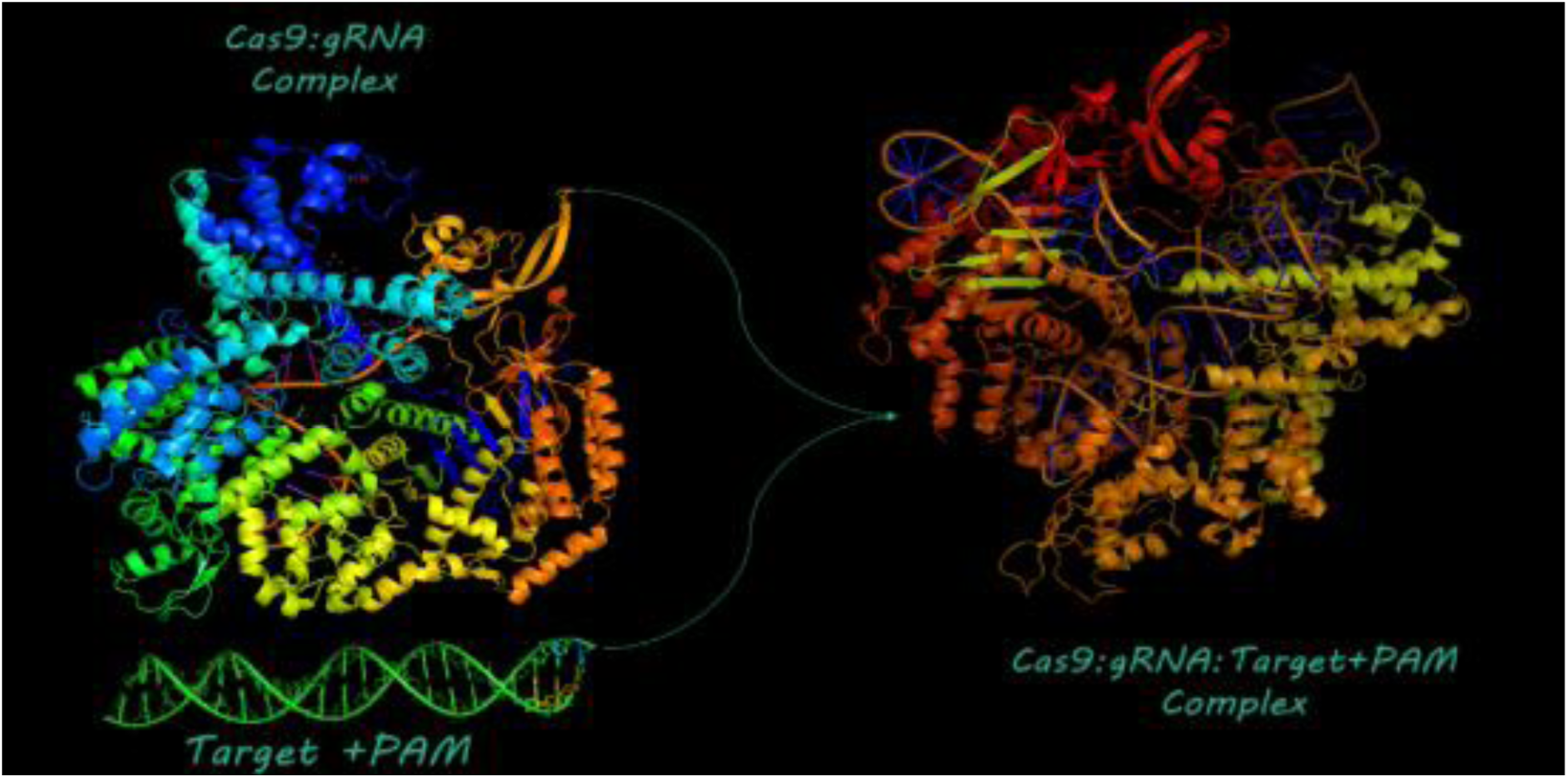
PyMol visualization for cas9: gRNA docked to Target DNA to identify cas9 cleavage of target DNA

#### Network Modelling Results

The ceRNA network and its graphical representation helped us to interpret the interaction mechanisms and signalling of circRNAs, miRNAs, and regulated mRNAs that have significant association to circRNAs regulatory role in HCC.

**Figure-22.**
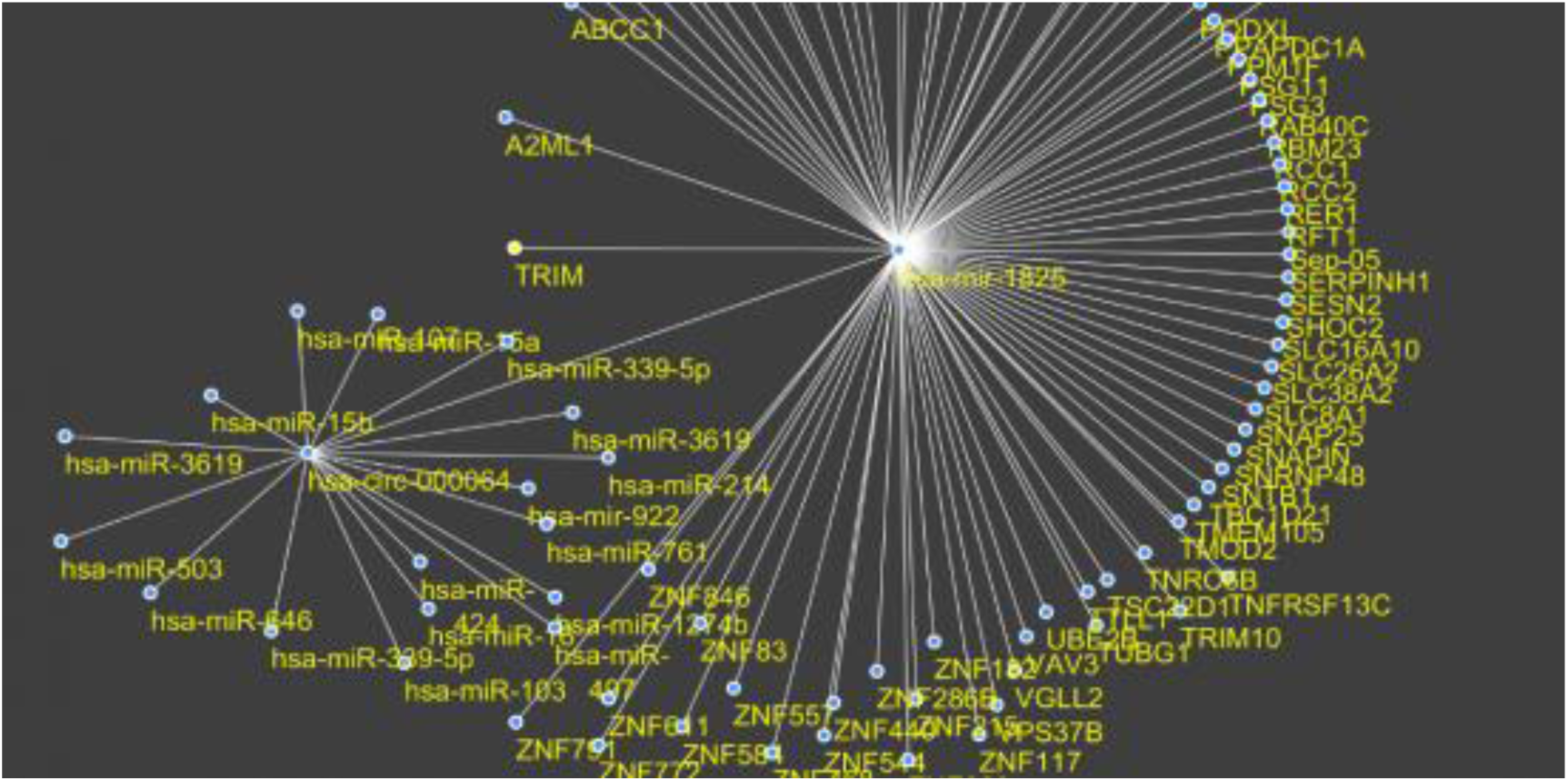
Regulatory network of circular RNA ceRNA network was constructed using Cytoscape 3.5.1 [43]. Network analysis mir-1825 targets network and circ-000064 associated miRNAs suggested the TRIM2 tumor suppressor which we used to experimentally assess the regulatory function of ceRNA network.

### Wet Lab Results

#### Gel electrophoresis of amplified Fragments

**Figure-23.**
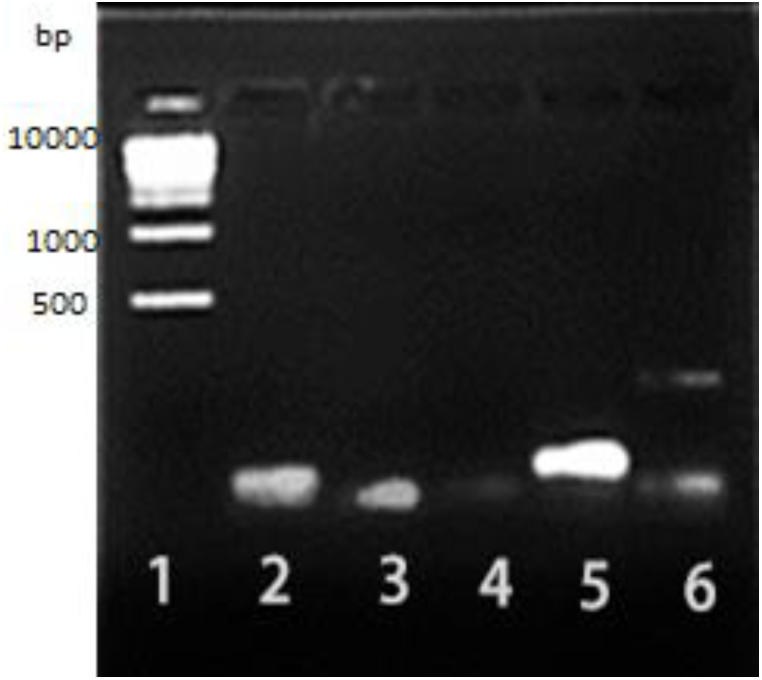
Agarose gel electrophoresis of amplified Fragments of first genetic construct (non-CRISPR-based): Lane 1: MW marker 500bp-10.000bp), Lane 2: Fragment 1 (CMV promoter +CMV enhancer), Lane 3: Fragment 2 (Laci+miRNA binding site+SVpoly A), Lane 4: Fragment 3(Lac promoter +Lac o+YFP+ Poly A signal), Lane 5: Fragment 4(cag promoter) at 1108bp, Lane 6: Fragment 5(hsa_circ_0000064+ terminator).

**Figure-24.**
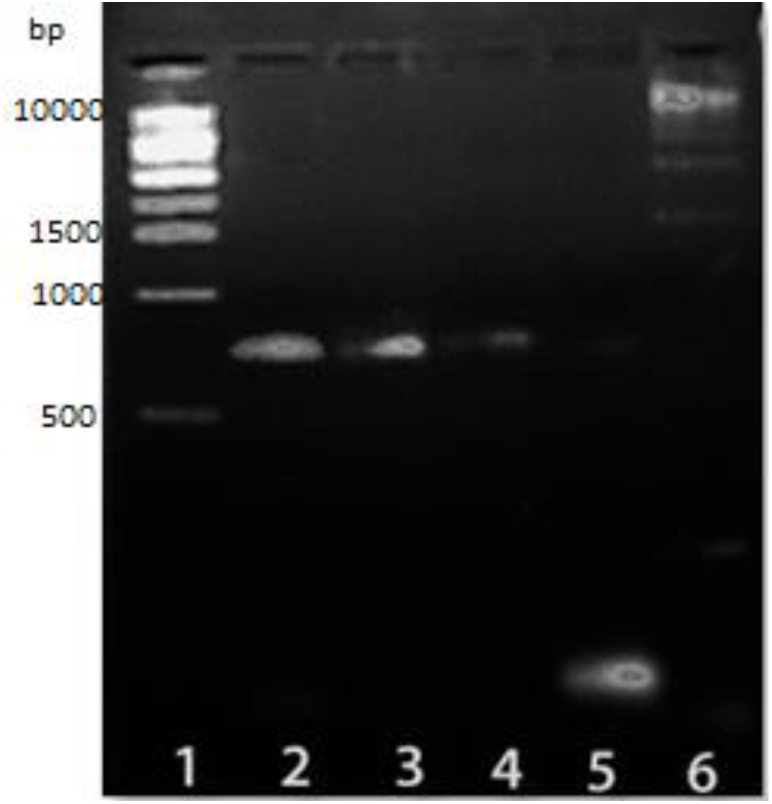
Agarose gel electrophoresis of amplified Fragments of second genetic construct (CRISPR-based): Lane 1: MW marker 500bp-10.000bp), Lane 2: Fragment 1 (U6 promoter + gRNA Target + gRNA scaffold + SV40 Terminator), Lane 3: Fragment 2(2[CMV enhancer CMV promoter + T7 promoter), Lane 4 Fragment 3(cas9; BBa_K1218011), Lane 5 Fragment 4(CYC1 Terminator), Lane 6: MW marker 500bp-10.000bp).

#### Competent E coli Transformation

**Figure-25.**
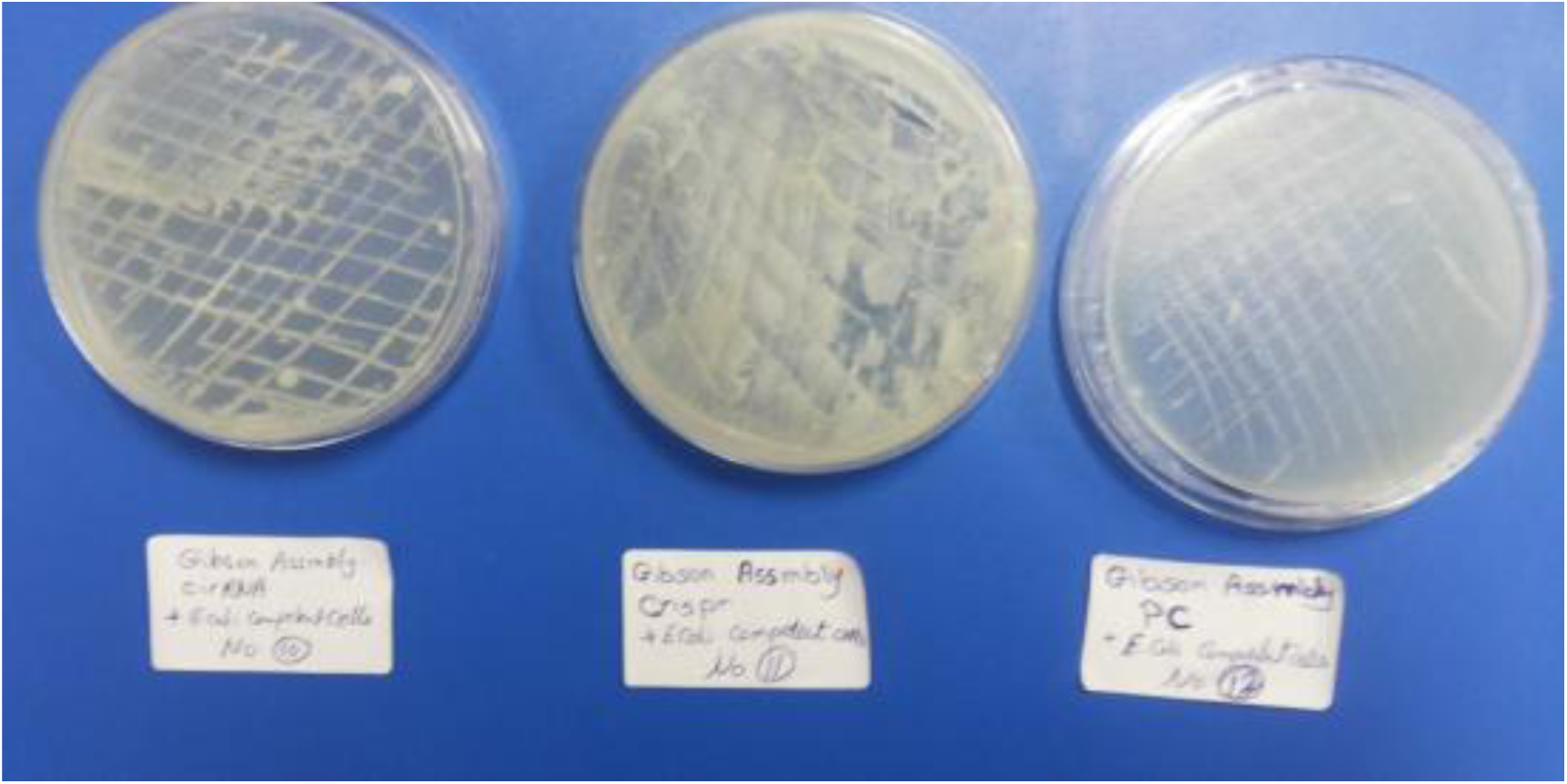
(Left to right) the first plate represents E-coli colony growth (+ve plasmid expressing hsa_circ_0000064, so E.coli can grow in LB with ampicillin due to AMPR gene. The second plate represents Marked E-coli colony growth (+ve plasmid expressing cas9 and vector carrying hsa_circ_0000064 and homology directed arm, so E.coli can grow in LB with ampicillin due to AMPR gene. The third plate represents E-coli colony growth (-ve plasmid) so E.coli cannot grow in LB media which contain ampicillin.

#### HepG2 transfection with CRISPR and non-CRISPR-based genetic construct or empty vector

**Figure-26.**
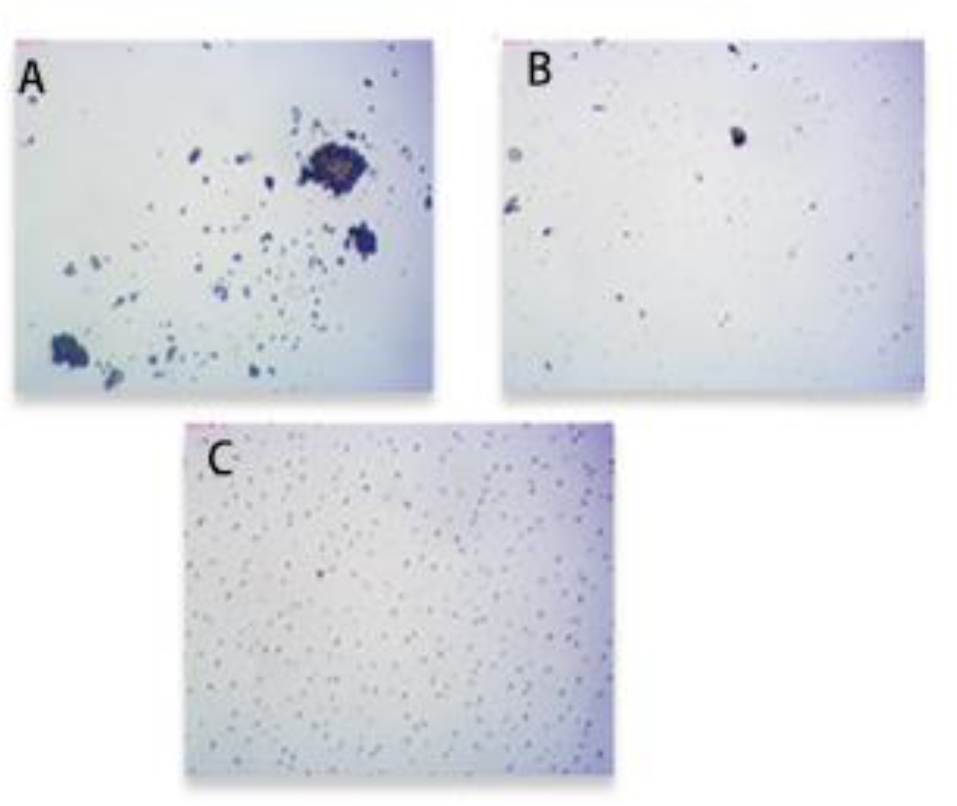
Assessment of transfection efficiencies. Cells were visualized 48 hours post-transfection under phase contrast microscope (A): Negative control (transfection with empty vector); (B): HepG2 transfected with genetic construct expressing hsa_circ_0000064: Shows decrease in cell count: Exogenous expression of hsa-circ-0000064 induces apoptosis in HepG2 cells. Phase contrast of the HepG2 cells two days after transient transfection with hsa-circ-0000064 expressing vector; (C):HepG2 transfected with genetic construct expressing Cas9 and pFETCh_vector expressing hsa_circ_0000064 and HDR shows marked apoptosis.

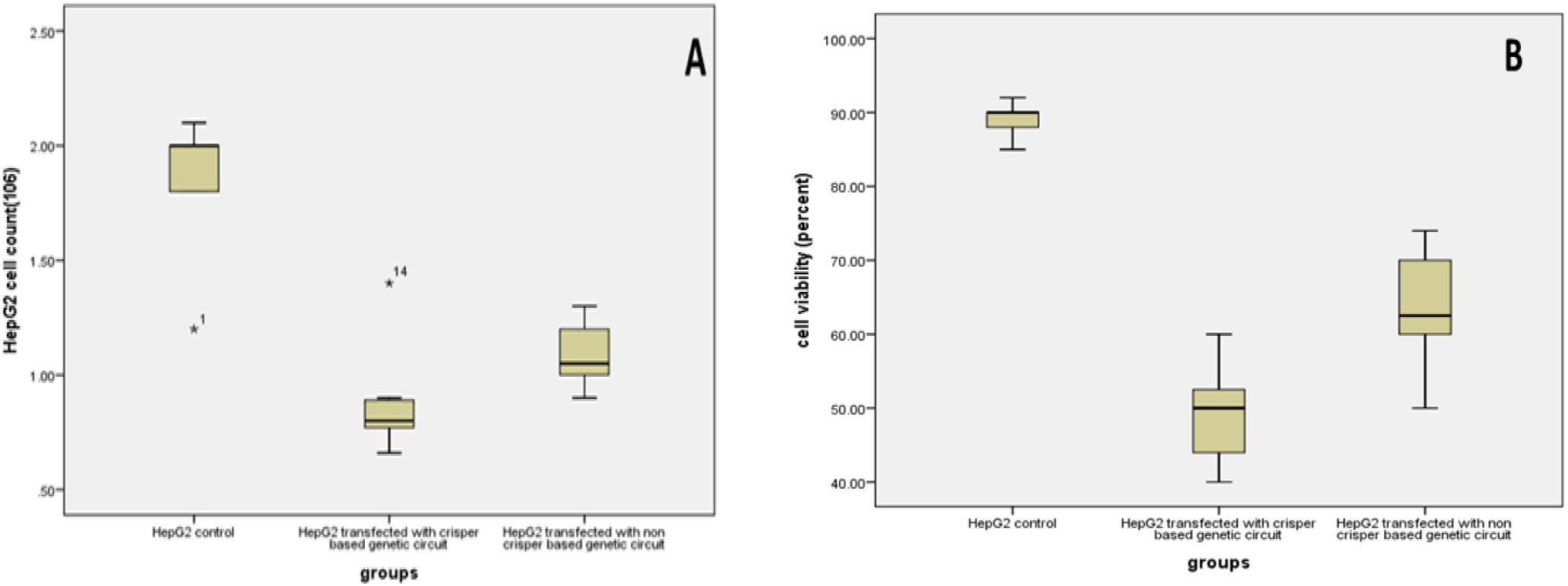

48 h after transfection, marked decreases in HepG2 cell counts and viability were observed by inverted microscopy. All the experimental conditions induced a cell-death phenotype that could be easily distinguished from control, indicating cell death after transfection with hsa-circ-0000064 via CRISPR and non-CRISPR-based genetic constructs with marked efficacy of CRISPR based genetic construct.

#### Expression of hsa_circ_0000064-miR-1285-TRIM2 mRNA in HEPG2 cell line after transient expression of hsa_circ_00064 via CRISPR and non-CRISPR-based genetic constructs

Our results showed that the up-regulation of *hsa_circ_0000064*, after transfection, was significantly associated with inhibition of miR-1285 expression (about 4 folds less) and up-regulation of *TRIM2 mRNA* which in turn inversely correlated with the cellular viability and the number of living HepG2 cells. These results indicate that hsa_circ_00064 may mediate TRIM2 up-regulation and subsequently show toxic effect in HCC cells. The present study showed that the CRISPR-based genetic circuit showed a markedly significant increase in *hsa_circ_0000064 expression than that induced by non-CRISPR-based method*.

**Figure-27.**
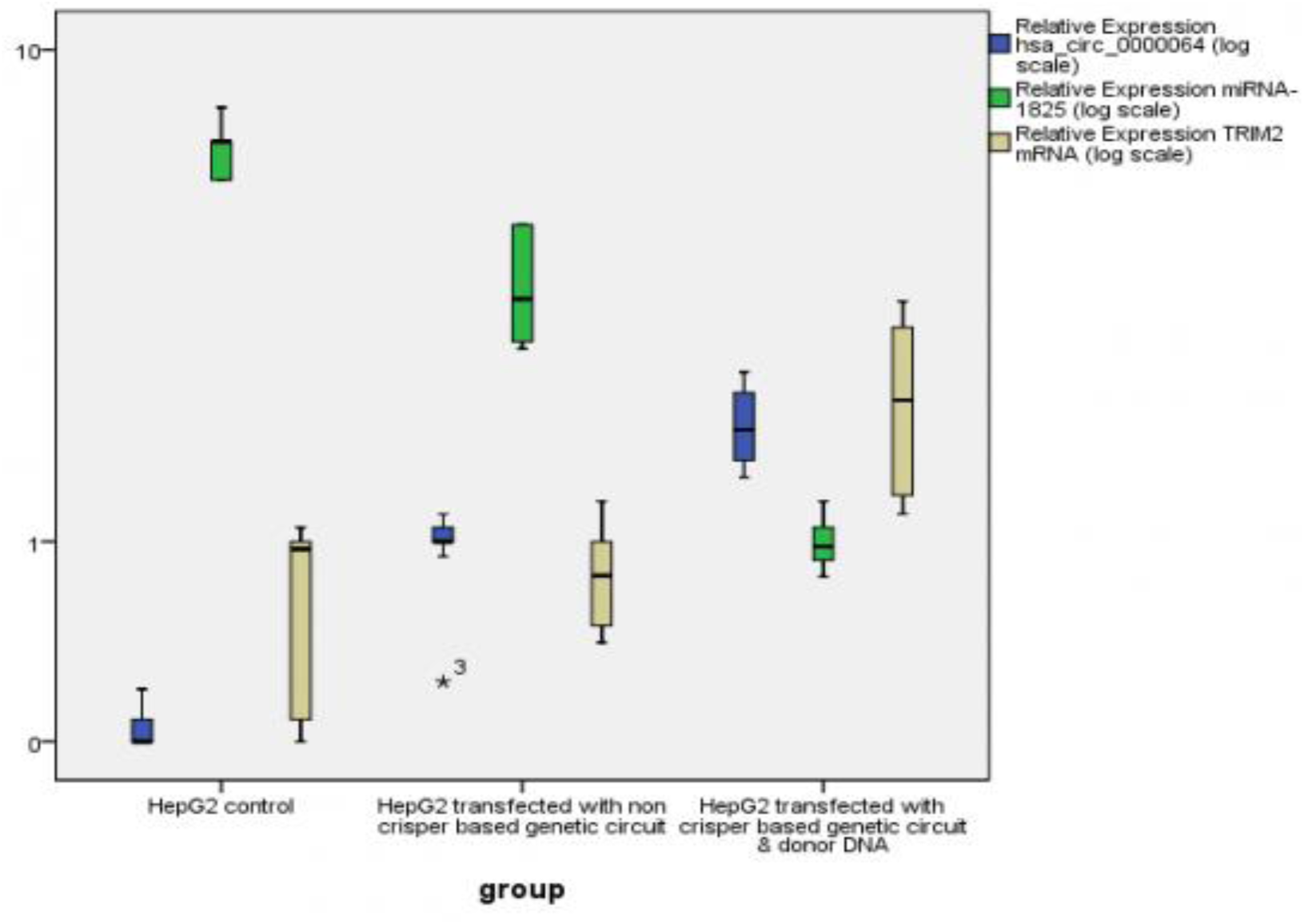
Box plot analysis representing the *effect of* HepG2 cell transfection with CRISPR-and non-CRISPR-based genetic constructs or a control *on the relative expression of the chosen genes by qPCR. (A) Hsa_circ_0000064; (B) miR-1285-3p; (C) TRIM2mRNA. The* results are expressed as the means±SD. *P<0.05

#### Flow Cytometery

It seems that has-circ-0000064 overexpression induces apoptosis via acting as sponge to miR-1285, resulting in up regulation of TRIM2 protein. TRIM2 functions as an E3 ubiquitin ligase that directs proteasome-mediated degradation of target proteins. Thus, it controls the apoptotic response acting as tumor suppressor gene.

**Figure-28.**
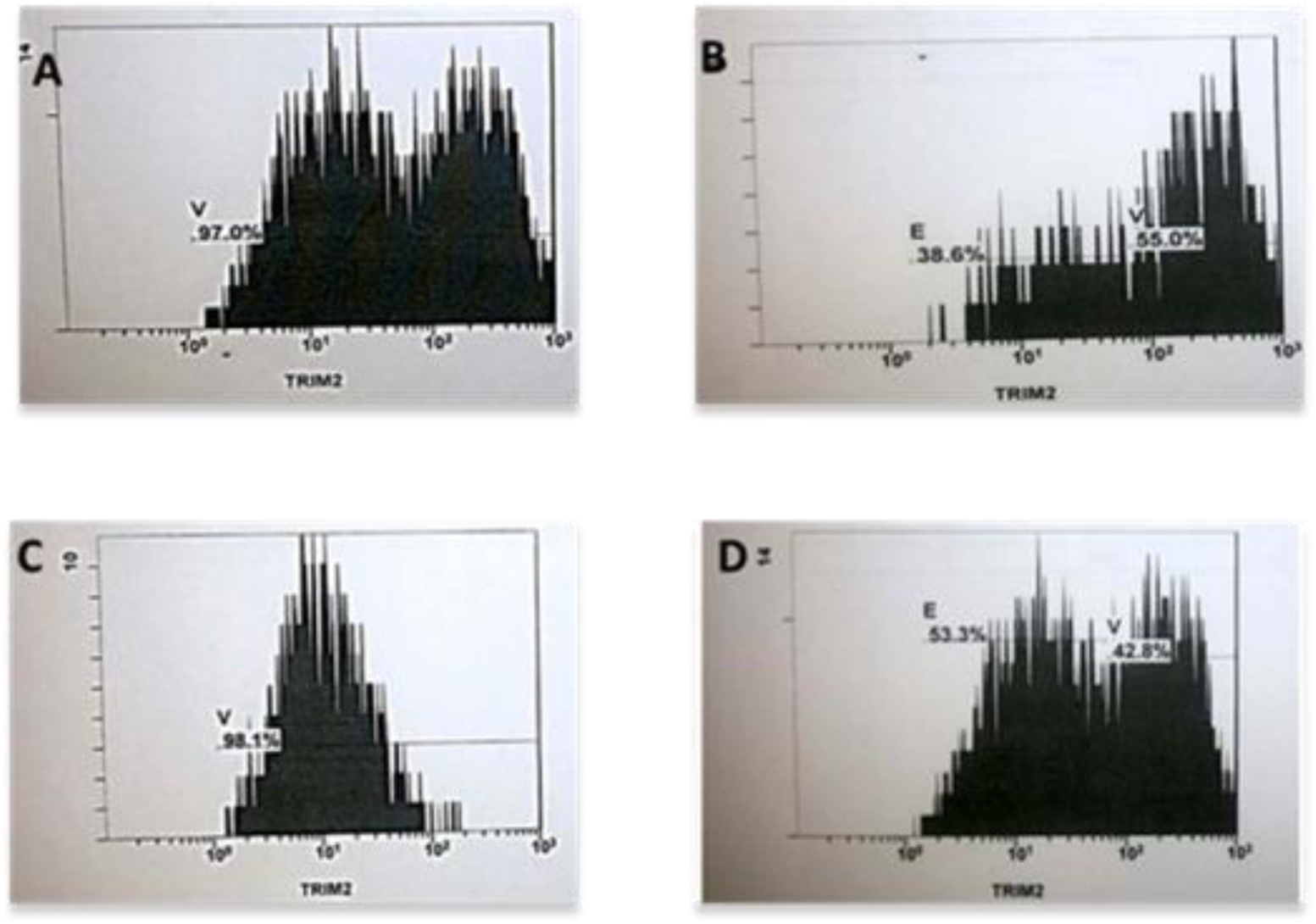
Flow cytometric analysis of HepG2 after transient transfection with hsa-circ-0000064 expressing vector or backbone. The cells overexpressing hsa-circ-0000064 showed accumulation of TRIM2 protein with mock. A: HepG2 transfected with CRISPR-based genetic construct; B: HepG2 transfected with non-CRISPR based genetic construct; C: Negative control; D: Positive control.

#### Immunohistochemical staining

Representative result of IHC staining of TRIM2 protein after transient transfection of has-circ-0000064 expressing vector mock in HepG2 cells. D. IHC staining of HepG2 cells after co-expression of hsa-circ-0000064 expressing vector shows the brightness of TRIM2 protein prominently in the HepG2 cells transfected with either CRISPR-or non-CRISPR-based genetic construct.

**Figure-29.**
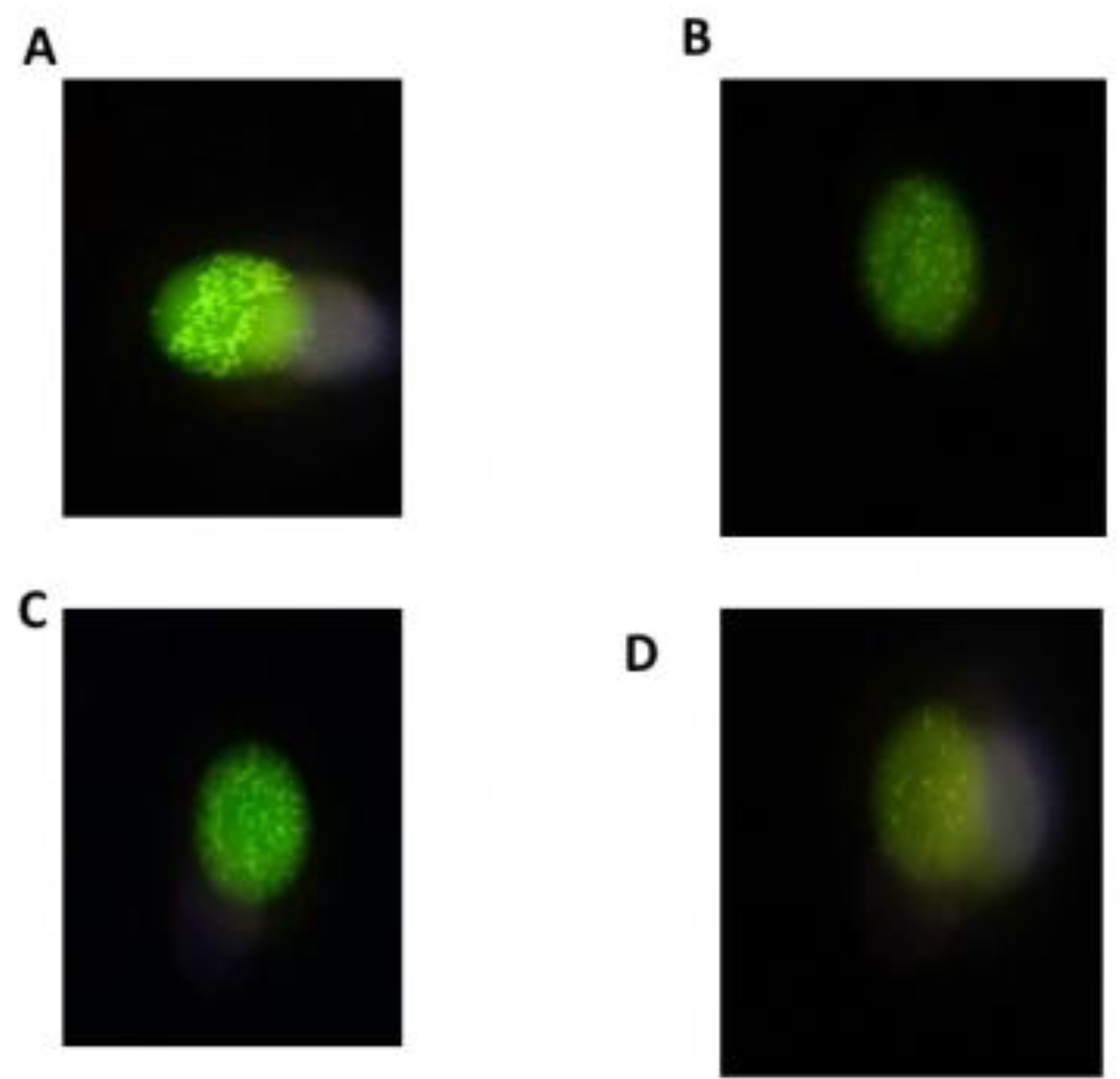
Immunohistochemical staining of HepG2 after transient transfection with hsa-circ-0000064 expressing vector or backbone. The cells overexpressing hsa-circ-0000064 showed accumulation of TRIM2 protein with mock. A: HepG2 transfected with CRISPR-based genetic construct; B: HepG2 transfected with non-CRISPR-based genetic construct; C: Positive control; D: Auto fluorescence.

## Discussion

Given the intricate interplay among the diverse RNA species, our team this year modified last year’s proposal with the application of CRISPR to knock-in our CircRNA gene into the genome of the HCC cell lines. RNA transcripts, like the long non-coding RNA and circular RNA, act as competing endogenous RNAs (ceRNAs) or natural microRNA sponges as they communicate with and co-regulate each other by competing for binding to shared microRNAs; a family of small non-coding RNAs that are important post-transcriptional regulators of gene expression. This regulation is scientifically effective way in manipulating critical roles in both normal physiology and tumorigenesis. miRNAs were revealed to repress their target genes via binding imperfectly to miRNA response elements (MREs) on the 3 ′ untranslated regions (3 ′ -UTRs) of target RNA transcripts and reducing expression of their target proteins either by mRNA breakdown or translational repression. Because each miRNA could target hundreds of genes and vice versa, each gene can be targeted by many miRNAs; such molecules are critically mentioned in the fine-tuned regulation of gene expression. RNAs functioning as in this course are named ceRNA. CeRNAs having common MREs can compete for binding of miRNA. It was suggested that these ceRNAs can talk to each other via their ability to compete for binding of miRNA. This cross-talk produces comprehensive cis and Trans organizing communication across all the transcriptome. Moreover, ceRNA networks further depend on the subcellular dispersion and tissue particularity of RNAs and miRNAs found in a specific cell type at a specific. The concentration of miRNAs is an important factor for ceRNA activity. If there are a less number of miRNAs than their targets, the ceRNA activity is reduced as the targets will remain largely unrepressed. Also, if there are more miRNAs as compared to their targets, there would have been no cross-regulation due to almost a universal repression of the targets regulating gene expression. Most circRNAs have been demonstrated as being tissue specific [10]. An example of the effect of circRNA-miRNA interaction is a study that evaluated the expression profile of human circRNAs in HCC tissues and identified circMTO1 (mitochondrial translation optimization homologue; hsa_circRNA_0007874/hsa_circRNA_104135) as being significantly down-regulated in HCC tissues. Bad prognosis and survival rate was detected in patients with low circMTO1. Via a biotin-labeled circMTO1 probe to preform RNA in vivo precipitation in HCC cells. Dan Han et al. identified miR-9 as the circMTO1-associated miRNA. It was also proven that circMTO1 suppresses progression of HCC by sponging miR-9 and thereby promoting p21 expression [47]. Acting in a ceRNA manner, circular RNA has_circ_0001564 has been proved to regulate mir-29c-3p in a sponge manner, which have significant role for regulating tumorigenicity in osteosarcoma [48]. Has_circ_0010729 have been shown to act through a ceRNA that includes mir-186/HIF-1α axis regulating apoptosis and proliferation of endothelial cells [49].

The association between circRNAs and cancer has been indicated in several recent studies mainly because of the major role that miRNAs play in gene regulation and cancer development. Therefore, there is a wide field of research open to new discoveries and validation of circRNA–miRNA–mRNA pathways, mainly related to cancer diagnosis and therapeutic treatments. This project provides a comprehensive comparison of genetic regulation effectiveness between competing endogenous RNA networks ceRNA versus CRISPR based genetic repair circuits.

## Conclusion

In this study, we adopted an integrative approach combining *in silico* data analysis with experimental validation. For the first time, we examined a set of HCC-related genes, cirRNA-miR-mRNA, linked to apoptosis. Based on the aforementioned findings, a reasonable interpretation of our hypothesis is that *hsa_circ_0000064* competes with *miR-1285 modulate the expression of TRIM2 mRNA that is closely linked to apoptosis pathway in liver. Interestingly, introducing this hsa_circ_0000064 into HCC cell line by CRISPR-based method seems to be more efficient than simply introducing plasmid transiently expressing hsa_circ_0000064 that may be promising therapeutic strategy for HCC*.

## Conflict of interest

The authors have declared no competing interest exist

