## Supplementary Materials for "Competing endogenous RNA network: Potential entrants to gene editing in Hepatocellular Carcinoma Gene editing of ceRNA in Hepatocellular Carcinoma"

**Response to Reviewers**

lgallagher: Hi AFCM Egypt, Please could you add a Conflict of Interest statement. A standard one for this, where no competing interests exist, would be: "The authors have declared that no competing interests exist."

**Response: We have added the conflict of interest statement.**

lgallagher: Hi AFCM Egypt, Please could you add a Financial Disclosure statement. In this, you should detail any funding received, who the funders were and you should explain any role the funders had in the study, ie: "The funders had no role in study design, data collection and analysis, decision to publish, or preparation of the manuscript." If no funding was recieved this should read: "The authors received no specific funding."

**Response: We have added the financial disclosure statement.**

gallagher: Hi AFCM Egypt, Please could you add a Data Availability statement? This should describe how readers and other researchers can access the minimal dataset needed to replicate the study. For a great example, check out University of Glasgow's statement here: https://synbioplos.breezio.com/article/4911588452237051730/campylocator-characterizing-genetic-circuit-components-in-e-coli-towards-a-campylobacter-jejunibiosensor

**Response: We have added the data availability statement.**

lgallagher: Hi AFCM Egypt, Thanks for your submission. We'd suggest establishing your research question early on in the Abstract.

**Response: We have included our simplified statement of our project aim to the background part of the abstract.**

aaa_2018: Avoid the ampersand (&) as a substitute for "and" in the main text. I'd leave out the word "special" to describe CRISPR - it's unclear what this means. Change the tense to past tense when describing research already carried out, eg: "[...] we used to induce forced [...]"

**Response: We have targeted these typographical errors, (&) was replaced with (and), the word (special) was omitted and the writing tenses were adjusted.**

lgallagher: Hi, just a copy-editing note: when you go back over your work check the punctuation carefully. There are some instances of extra spaces between words or around punctuation, and although these don't affect the scientific meaning of the paper they can be a little distracting for readers.

**Response: We managed to review our text in terms of punctuation and extra-spaces were removed.**

PreethaG: The conclusion section in your abstract doesn't need to summarize your experimental approach, and could be made more concise if that summary was omitted.

**Response: We modified the conclusion section in the abstract to make it more concise.**

lgallagher: Hi, it's great that you cite a high number of related studies and existing literature as it helps to put your work into perspective. Just a quick copy-editing note: when you write references try to do these consistently ie: [1], [1,2,3] etc. this just makes for slightly easier reading and means that these will link to the reference section so that they can be easily checked or located.

**Response: We have rewritten the citation notations to be in line with the text and easily checked.**

Dario: Hi, I'd suggest adding in the introduction some specific references that support the ceRNA role of circRNAs and the role of circRNAs in HCC. Also, it would be worth clarifying in the text that circRNAs can also act as ceRNAs

**Response: We have added more specific references in both introduction and discussion sections with clarification of circular RNA role as ceRNA.**

aaa_2018: Minor comment: One of the formulas is vertically compressed in this image.

**Response: We have adjusted the equation figures to avoid this vertical compression.**

aaa_2018: I'd put this reference in the legend, since it's the same for every parameter.

**Response: We have readjusted the reference for parameters.**

aaa_2018: (Same comment about the reference, since it's the same for all parameters)

**Response: We have readjusted the reference for parameters.**

aaa_2018: Do the colours on this figure mean anything?

**Response: We have detailed the indications for colors through the figure caption.**

PreethaG: Figure captions should not be bolded. (I noticed this happen quite a few times throughout your paper -- it would be a good idea to check all captions).

**Response: We have rechecked the captions.**

PreethaG: This figure caption should not be bolded and should probably be more descriptive (include your modelling approach + software and a brief description of the modelling results).

**Response: We have readjusted the caption and added more description about the modelling approach, software and modelling results.**

aaa_2018: You could label the DNA size marker ladder to give an idea of scale.

**Response: DNA ladder has been labeled.**

aaa_2018: (Same comment about the size marker lane)

**Response: DNA ladder has been labeled.**

aaa_2018: This might be just a display problem with this problem, but this figure is a bit hard to read.

**Response: This figure has been adjusted to be easily viewed.**

aaa_2018: The sentence "Well what is the story?" is a bit informal for a scientific article. I'd recommend something like: "There are different possible explanations for what we have observed." The last sentence also seems disconnected: "Such competing endogenous RNAs."

**Response: Informal sentences and incomplete sentences have been reviewed.**

aaa_2018: I think the last word of the article has been cut off.

**Response: Text consistency has been reviewed and adjusted.**

lgallagher: Hi AFCM Egypt, Have you considered showing your flow cytometry data in the form of dot-plots rather than histograms? We'd also suggest that you make your research question clear from the start and describe your methods in a way that is easily reproducable.

**Response: We think it might be technically difficult to get dot-plots instead of histograms as the original platform at the time of the experiment didn't offer these plots. Consequently we managed to make our project aim clearer. Methods were also edited to make the project more reproducible.**

aaa_2018: CRISPR is a very hot topic at the moment!

**Response: Thanks for your review, we hope to provide promising pipeline for CRISPR editing of ceRNA netoworks.**
